# A redundant chloroplast protein is co-opted by potyvirids as the scaffold protein to mediate viral intercellular movement complex assembly

**DOI:** 10.1101/2023.09.29.560214

**Authors:** Li Qin, Hongjun Liu, Peilan Liu, Lu Jiang, Xiaofei Cheng, Fangfang Li, Wentao Shen, Zhaoji Dai, Hongguang Cui

## Abstract

For viruses in the family *Potyviridae* (potyvirids), three virus-encoded proteins (P3N-PIPO, CI and CP) and several host components are known to coordinately regulate viral cell-to-cell movement. Here, we found that HCPro2 encoded by areca palm necrotic ring spot virus is involved in the intercellular movement, which could be functionally complemented by its counterpart HCPro from a potyvirus. The affinity purification and mass spectrum analysis identified several viral factors (including CI and CP) and a variety of host proteins that physically associate with HCPro2. We demonstrated that HCPro2 interacts with either CI or CP *in planta*, and the three form plasmodesmata (PD)-localized interactive complex in viral infection. Further, we screened HCPro2-associating host proteins, and identified a common host protein RbCS that mediates the interactions of HCPro2-CI, HCPro2-CP and CI-CP among the complex. Knockdown of *NbRbCS* simultaneously impairs the interactions of HCPro2-CI, HCPro2-CP and CI-CP, and significantly attenuates the intercellular movement and systemic infection for ANRSV and other three tested potyvirids. This study highlights that a nucleus-encoded chloroplast-targeted protein is hijacked by potyvirids as the scaffold protein to mediate the assembly of viral intercellular movement complex to promote viral infection.

## Introduction

Plasmodesmata (PD) are plasma-membrane-lined nanochannels that cross rigid cell wall between adjacent cells in higher plants, allowing the exchange of signals and resources among cells for developmental regulation and stress responses (Lucas et al., 2009; Faulkner, 2018; Sager & Lee, 2018; Li et al., 2023). Plant viruses, as the obligate intracellular parasites, take full advantage of PD to spread intercellularly to establish systemic infection. However, the small aperture of PD allows small molecules to diffuse between cells, but physically restricts the passage of macromolecules or macromolecular complexes such as viral ribonucleoprotein complexes (vRNPs) or virions (Oparka, 2004; Lucas et al., 2009; Wang, 2021). To overcome this barrier, plant viruses evolved diverse types of movement proteins (MPs) that interact with host proteins to modify PD to translocate vRNPs or virions between adjoining cells (Harries & Ding, 2011; Ueki & Citovsky, 2011; Vijayapalani et al., 2012; Navarro et al., 2019; Wang et al., 2021). Currently, three are three modes assigned to the cell-to-cell movement of plant viruses (Wu & Cheng, 2020; Wang, 2021): i) Single MP encoded by tobacco mosaic virus and its related viruses targets PD to increase the size exclusion limit (SEL) of PD for the trafficking of vRNPs or virions (Kawakami et al., 2004; Wright et al., 2007; Ueki & Citovsky, 2011; Butkovic et al., 2023); ii) Triple-gene-block viruses such as potexviruses and hordeiviruses encode three MPs that coordinately transport vRNPs or virions through PD (Verchot-Lubicz, 2005; Jackson et al., 2009; Lim et al., 2009; Verchot-Lubicz et al., 2010; Verchot., 2022); iii) The MPs encoded by viruses in three families *Secoviridae*, *Bromoviridae* and *Caulimoviridae* form tubule structures that traverse through PD for virions movement (Laporte et al., 2003; Pouwels et al., 2004). However, the cell-to-cell movement for viruses in the family *Potyviridae* (potyvirids), the largest group of plant-infecting RNA viruses, has not been categorized, and is still an understudied area (Wang, 2021).

All potyvirids excluding bymoviruses possess one positive-sense, single-stranded RNA genome (∼ 9.7 kb), which contains a long, full-genome open reading frame (ORF) and a relatively short ORF (PIPO) embedded in P3-coding region (Chung et al., 2008; Inoue-Nagata et al., 2022). PIPO becomes translatable in frame with P1-to-P3N-coding region from viral genomic subpopulation, which origin from viral RNA polymerase (NIb) slippage during viral replication (Olspert et al., 2015; Rodamilans et al., 2015; Mingot et al., 2016; Untiveros et al., 2016; Yang et al., 2021). Two different polyproteins after translation are proteolytically processed by virus-encoded proteases into 10 to 12 mature units (Cui & Wang, 2019; Yang et al., 2021). None of these viral factors is indicated as viral MP, but three of them, P3N-PIPO (a translational fusion of N-terminus of P3 with PIPO), CI (cylindrical inclusion protein) and CP (coat protein), are known to regulate viral intercellular trafficking in a coordinated manner.

P3N-PIPO is a PD-localized viral factor, facilitating its own cell-to-cell movement (Wei & Wang, 2008; Vijayapalani et al., 2012; Cheng et al., 2017). Disrupting the generation of P3N-PIPO in different potyvirids restricts viral cell-to-cell movement but does not affect viral replication (Choi et al., 2005; Wen et al., 2010; Geng et al., 2015; Cui et al., 2017). CI, an RNA helicase required for viral genome replication, is a multifunctional protein (Sorel et al., 2014). Accumulated genetic evidences assign an independent role for CI in viral intercellular movement (Carrington et al., 1998; Sorel et al., 2014; Deng et al., 2015). CI is recruited to PD via an interaction with P3N-PIPO, and forms conical structures that anchor to and extend through PD (Wei et al., 2010; Cui et al., 2017; Wang, 2021). Cytopathological and biochemical evidences revealed that CP or virion binds with CI-forming conical structures at PD to aid viral intercellular passage (Rodriguez-Cerezo et al., 1997; Roberts et al., 1998; Gabrenaite-Verkhovskaya et al., 2008). Mutating of CP that disrupts viral particle assembly compromises the intercellular spread as well (Dolja et al., 1994; Dolja et al., 1995; Seo et al., 2013; Dai et al., 2020), suggesting that viral cell-to-cell movement occurs likely in the form of virion (Wang et al., 2021). Helper component-protease (HCPro) is another multitasking protein, and its function in RNA silencing suppression (RSS) is well-studied (Valli et al., 2018; Cui & Wang, 2019; Pasin et al., 2022). Based on previous observations, HCPro might participate in viral cell-to-cell movement: i) HCPro is able to traffick from cell to cell, increase the SEL of PD, and facilitate the intercellular movement of viral RNA (Rojas et al., 1997); ii) HCPro stabilizes CP and enhances the yield of virions (Valli et al., 2014), thereby promoting viral intercellular movement likely in an indirect manner (Wang, 2021); iii) HCPro or CI could form a protrusion at one end of virion, in the case of potato virus A (PVA) (Torrance et al., 2006; Gabrenaite-Verkhovskaya et al., 2008). Nevertheless, a detailed survey on the interactions of HCPro with PD-localized movement-related proteins needs to be performed to clarify its role(s) in viral intercellular movement.

The cell-to-cell movement of plant viruses usually depends on the coordinated action of viral MPs and host factors (Schoelz et al., 2011; Heinlein, 2015; Wang, 2015; Reagan & Burch-Smith, 2020). A few host proteins have been identified to interact with potyvirid MPs, facilitating viral intercellular trafficking. A hydrophilic plasma membrane-associated cation-binding protein in *Arabidopsis* (AtPCaP1) is recruited to PD via its interaction with P3N-PIPO to promote the cell-to-cell movement of turnip mosaic virus (TuMV, *Potyvirus*) (Vijayapalani et al., 2012; Cheng et al., 2020). In parallel, an AtPCaP1 homolog in *Nicotiana benthamiana*, called as NbDREPP, contributes to the intercellular movement of tobacco vein banding mosaic potyvirus (Geng et al., 2015). PCaP1 might function in the anchoring of P3N-PIPO to PD (Vijayapalani et al., 2012), and serves actin filaments in PD to increase the SEL of PD (Cheng et al., 2020). Another plasma membrane protein - synaptotagmin A positively regulates the intercellular trafficking of TuMV P3N-PIPO through PD (Uchiyama et al., 2014). An α-expansin in *N. benthamiana* - NbEXPA1 promotes either the replication or cell-to-cell spread of TuMV (Park et al., 2017).

It has long been known that chloroplast is involved in plant-virus interactions, and the subject of chloroplast-virus interplay attracts great interest and more attention (Li et al., 2015; Zhao et al., 2016; Bhattacharyya & Chakraborty, 2018). An increasing number of nucleus-encoded chloroplast-targeted proteins are co-opted by different plant viruses for viral replication, movement or/and counteracting host defense response (Medina-Puche et al., 2020; Cheng et al., 2021; Ji et al., 2021; Chen et al., 2022; Han et al., 2023). During photosynthetic pathway, ribulose 1, 5-bisphosphate carboxylase/oxygenase (Rubisco) is responsible for catalyzing the first rate-limiting step in CO_2_ fixation. Rubisco is comprised of eight large subunits (RbCL; ∼50-55 kDa) and eight small subunits (RbCS; ∼12-18 kDa) which form a hexadecameric L_8_S_8_ complex (Bracher et al., 2017; Mao et al., 2023; Prywes et al., 2023). RbCL is encoded by chloroplast genome, whereas RbCS is nucleus-encoded (Bracher et al., 2017). RbCS interacts with tobamoviral MPs at PD for intercellular and long-distance movement (Zhao et al., 2013). Both RbCL and RbCS interact with either P3 or P3N-PIPO in cases of several viruses in *Potyvirus* genus (potyviruses) (Lin et al., 2011). In addition, RbCL interacts with HCPro of bean common mosaic potyvirus, and CP of potato virus Y (Feki et al., 2005; Kumar et al., 2020). Anyway, the biological relevance underpinning these interactions is needed to be investigated.

Previously, we characterized two novel viruses - areca palm necrotic spindle-spot virus (ANSSV) and areca palm ring spot virus (ANRSV) belonging into a new genus *Arepavirus* of *Potyviridae* family (Yang et al., 2018; Cui & Wang, 2019; Yang et al., 2019). Both viruses exhibit a distinct pattern of leader proteases – HCPro1 and HCPro2 in tandem, which are short in size and share low sequence similarity with their counterparts in other potyvirids (Cui & Wang, 2019; Qin et al., 2021). This promoted us to investigate the functions of HCPro1 and HCPro2 during viral infection. We found that HCPro1 is dispensable for viral infection, whereas HCPro2 not. Besides acting as the viral suppressor of RNA silencing (VSR), HCPro2 is involved in the cell-to-cell movement, which could be functionally complemented by its homolog from a potyvirus. We demonstrated the three viral factors (HCPro2, CI and CP) form interactive complex at PD in the presence of viral infection. We further identified a common host protein - NbRbCS that mediates the interactions among them. Knockdown of *NbRbCS* greatly impairs viral intercellular movement and systemic infection for ANRSV and other three tested viruses in *Potyvirus* genus. Altogether, this study suggests that a redundant nucleus-encoded chloroplast-targeted protein is hijacked by potyvirids to mediate and reinforce the assembly of viral movement complex.

## Results

### HCPro1 is dispensable for ANRSV infection

Previously, we developed a GFP-tagged ANRSV clone - pRS-G (Wang, et al., 2021). To investigate the functional role(s) of HCPro1 during ANRSV infection, the entire HCPro1-coding region was removed from pRS-G to produce the clone pRS-G(△HCPro1) (Fig. 1A). pRS-G and pRS-G(△HCPro1) were each inoculated into ten *N. benthamiana* seedlings via agroinfiltration (OD_600_ = 0.5 per clone). At different time points, all plants inoculated with either pRS-G or pRS-G(△HCPro1) exhibited dwarfism and leaf rugosity symptoms (Fig. 1B), as well as obvious GFP signals along veins in top non-inoculated leaves (Fig. 1B). Interestingly, the more serve symptoms along with stronger fluorescence intensity were observed in plants inoculated with the HCPro1-deleted virus clone (Fig. 1B). Consistent with above observations, either GFP or viral genomic RNA was more accumulated in plants inoculated with pRS-G(△HCPro1) (Fig. 1C, D). Given that the deletion of HCPro1-coding sequence shortens the size of viral genome, it is uncertain whether the enhancement effect on viral infectivity is attributed to an alteration of viral genome, or a negatively regulatory role exerted by HCPro1 protein. Nevertheless, our current data support a notion that HCPro1 is dispensable for ANRSV infection in the model plant - *N. benthamiana*.

**Fig. 1.**
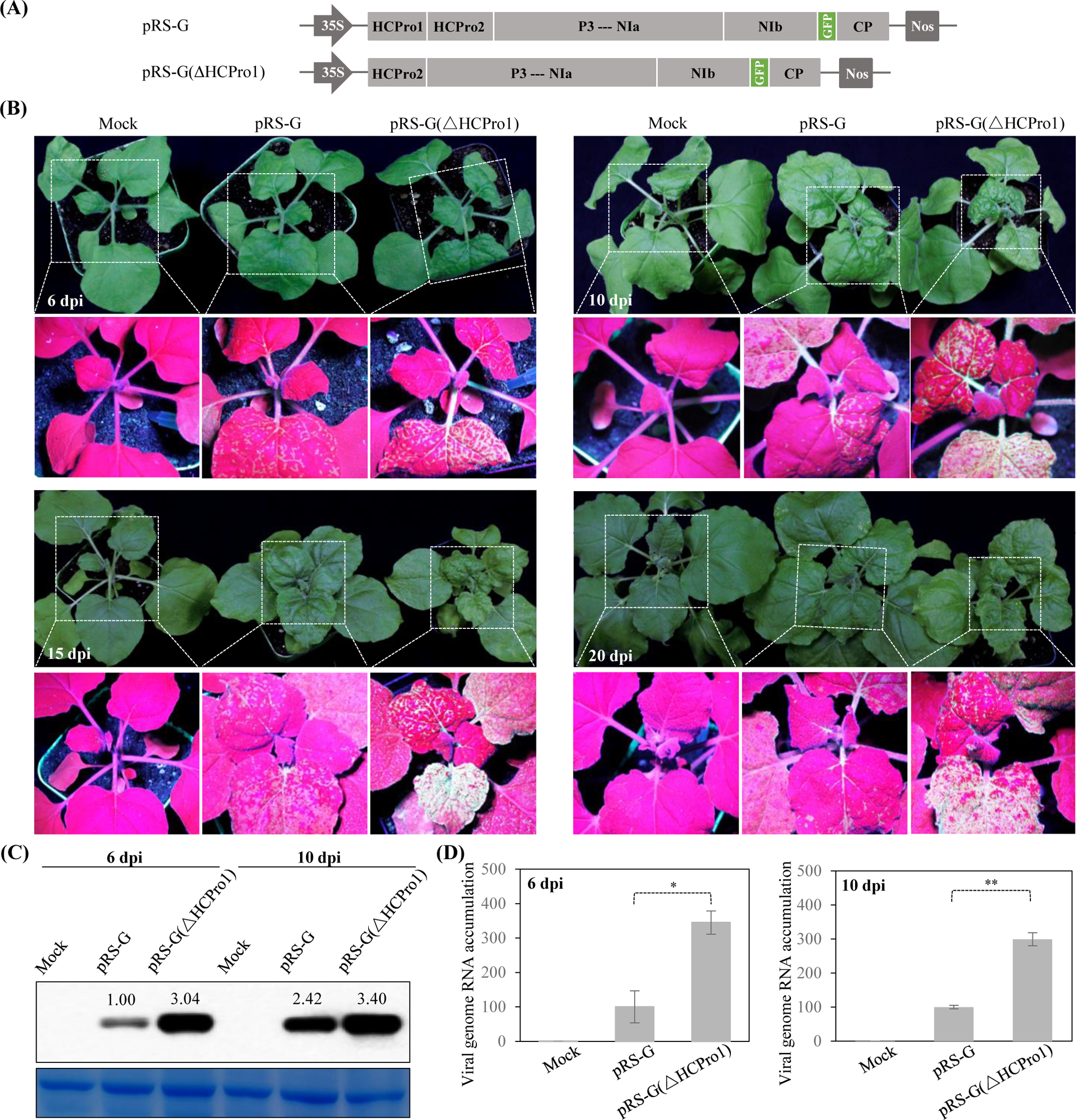
HCPro1 is dispensable for ANRSV infection. **(A)** Schematic diagrams of pRS-G and pRS-G(△HCPro1). P3---NIa the coding regions of seven viral factors, including P3, P3N-PIPO, 6K1, CI, 6K2, VPg and NIa-Pro. **(B)** Infectivity test of pRS-G and pRS-G(△HCPro1) in *N. benthamiana*. The representative *N. benthamiana* plants inoculated with the indicated virus clones were photographed under either daylight (upper) or UV light (lower). Mock, empty vector control. **(C)** Western blot analysis of GFP accumulation in inoculated *N. benthamiana* plants. Total proteins were extracted from top non-inoculated leaves at the indicated time points. The hybridization signal intensity was quantitatively analyzed with ImageJ software (Schneider et al., 2012). Coomassie blue staining of RbCL was used as a loading control. **(D)** Real-time qPCR analysis of viral RNA accumulation in inoculated plants. Total RNAs were extracted from top non-inoculated leaves, followed by real-time RT-qPCR analysis. The values represent the mean ± standard deviation (SD) from three independent biological replicates. The average values for pRS-G were designated as 100 to normalize the data. Statistically significant differences, determined by an unpaired two-tailed Student’s *t* test, are indicated by asterisks. *, 0.01<*P*<0.05; **, 0.001<*P*<0.01.

### HCPro2 functions in viral cell-to-cell movement, which could be functionally complemented by its counterpart - HCPro from a potyvirus

To examine the functions of HCPro2 during viral infection, we completely deleted HCPro2-coding sequence in pRS-G to generate the clone pRS-G(△HCPro2) (Fig. 2A). Infectivity test showed that all eight *N. benthamiana* plants infiltrated with pRS-G displayed obvious GFP fluorescence in upper non-inoculated leaves at 8 dpi and 16 dpi, whereas the eight plants inoculated with pRS-G(△HCPro2) not. RT-PCR assay confirmed that these plants infiltrated with pRS-G(△HCPro2) were not systemically infected. Previously, we demonstrated that ANSSV HCPro2 (ssHCPro2) expresses RSS activity (Qin, et al., 2021). Thus, ANRSV HCPro2 might execute RSS function as well, and its deletion would abolish viral infectivity. To test the RSS activity of ANRSV HCPro2, we constructed three T-DNA vectors for respective expression of HA-tagged HCPro1 (HCPro1-HA), HCPro2 (HCPro2-HA), and HCPro1-HCPro2 (HCPro1-HCPro2-HA) of ANRSV. Each of these constructs, together with a plasmid for expressing GFP reporter (Li et al., 2014) were co-inoculated into *N. benthamiana* leaves via agroinfiltration. Co-expression of GFP along with either empty vector or HA-tagged ssHCPro2 (ssHCPro2-HA) was included as negative and positive controls, respectively (Qin et al., 2021). At 60 hours post-inoculation (hpi), leaf patches co-expressing HCPro2-HA/GFP, HCPro1-HCPro2-HA/GFP or ssHCPro2-HA/GFP displayed strong GFP fluorescence, whereas no obvious fluorescence was observed on leaf patches co-expressing HCPro1-HA/GFP or negative control (Supplemental Fig. S1A). Consistent with above observations, a higher abundance of GFP at either protein or RNA level was observed in leaf patches co-expressing HCPro2-HA/GFP, HCPro1-HCPro2-HA/GFP or ssHCPro2-HA/GFP (Supplemental Fig. S1A, B), indicating that HCPro2 is the VSR protein of ANRSV.

**Fig. 2.**
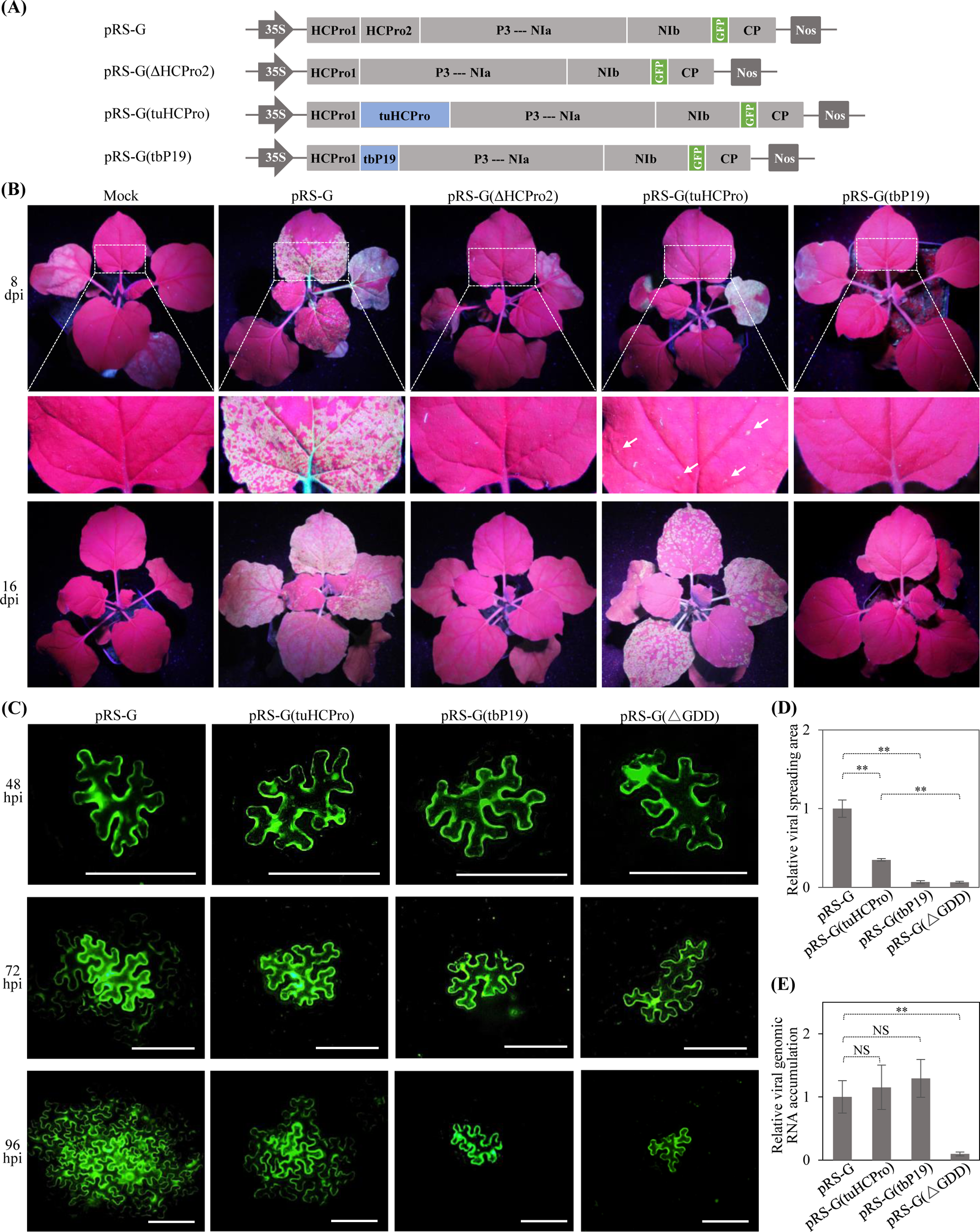
Effects of either deletion of HCPro2 or its substitution with different VSRs on viral infectivity. **(A)** Schematic diagrams of the derivatives of pRS-G. TuMV HCPro and TBSV P19 are represented by tuHCPro and tbP19, respectively. **(B)** Infectivity test of the derivatives of pRS-G in *N. benthamiana*. Representative photographs were taken under UV light at the indicated time points. The close view of leaf regions indicated by dashed boxes is shown. White arrows indicate fluorescence spots. Mock, empty vector control. **(C)** Time course observation of viral cell-to-cell movement for the indicated virus clones. Viral intercellular movement was monitored at 48 hpi, 72 hpi, and 96 hpi. Bars, 100 μm. **(D)** Statistical analysis of the size of viral spreading area at 96 hpi. At least 25 infection foci per clone from three independent experiments was analyzed. The size of infection foci is was calculated by ImageJ. The data are presented as the mean ± SD (*n* ≥ 25). Statistically significant differences, determined by an unpaired two-tailed Student’s *t* test, are indicated by asterisks. **, 0.001<*P*<0.01. **(E)** The effects of hybrid virus clones on viral genomic RNA accumulation. Relative viral genomic RNA accumulation was determined by real-time RT-qPCR. Error bars denote the SD from three biological replicates. **, 0.001<*P*<0.01; NS, no significant difference.

The potyvirus-encoded HC-Pro (the counterpart of ANRSV HCPro2), together with an unrelated P19 protein encoded by tombusvirus, are well-studied VSRs (Anandalakshmi et al., 1998; Kasschau & Carrington, 1998; Silhavy et al., 2002). To explore additional functions of HCPro2 beyond RSS, we substituted HCPro2 in pRS-G with either TuMV HCPro (tuHCPro) or P19 of tomato bushy stunt virus (tbP19) to produce two hybrid clones - pRS-G(tuHCPro) and pRS-G(tbP19) (Fig. 2A). *N. benthamiana* seedlings (*n* = 10 per clone) were inoculated with pCB301 (negative control), pRS-G (positive control), pRS-G(tuHCPro) and pRS-G(tbP19), followed by observations under ultraviolet (UV) light in every one- to two-day interval for one month. At 8 dpi, obvious fluorescence spots were observed in upper non-inoculated leaves of plants inoculated with pRS-G(tuHCPro). All plants inoculated with either pRS-G(tuHCPro) or wild-type pRS-G displayed comparable but not identical pattern of fluorescence signals at 13, 16 and 30 dpi (Fig. 2B; Supplemental Fig. S2A). In contrast, only three out of 10 plants infiltrated with pRS-G(tbP19) showed scattered fluorescence spots in only one non-inoculated leaf at 30 dpi (Supplemental Fig. S2A). Altogether, HCPro2 implements additional function(s) beyond RSS during viral infection, which could be largely complemented by its counterpart in TuMV.

Further, we examined the performance of hybrid viruses in intercellular movement. Agrobacterial cultures harboring pRS-G, pRS-G(tuHCPro), pRS-G(tbP19) and pRS-G(△GDD) (a replication- and movement-null mutant that lacks a strictly-conserved GDD motif in viral RNA polymerase) was highly diluted to 0.0001 of OD_600_, followed by infiltration into *N. benthamiana* leaves. At 48 hpi and 60 hpi, single cells emitting GFP fluorescence, representing primarily-transfected cells, were monitored for all clones (Fig. 2C; Supplemental Fig. S2B). Clear viral spreading from primarily-transfected to peripheral cells started at 72 hpi for pRS-G, and 84 hpi for pRS-G(tuHCPro) (Fig. 1C; Supplemental Fig. S2B). Thus, replacement of HCPro2 with tuHCPro partially inhibited viral intercellular spread (Fig. 2D; Supplemental Fig. S2C). In contrast, pRS-G(tbP19) was deficient in viral cell-to-cell movement, as in the case of pRS-G(△GDD) (Fig. 2C; Supplemental Fig. S2B). Moreover, we assessed the performance of hybrid viruses in viral genomic RNA accumulation. *N. benthamiana* leaves inoculated with these clones (OD_600_ = 0.3 per clone) were assayed by real-time RT-qPCR at 60 hpi as the viral intercellular movement did not occur at this time point (Supplemental Fig. S2B). As shown in Fig. 2E, no significant difference on viral genomic RNA accumulation was found between wild-type pRS-G and each of hybrid clones.

In conclusion, ANRSV HCPro2, besides acting as the VSR, also functions in viral cell-to-cell movement, and the function could be largely complemented by its counterpart-HCPro from TuMV.

### HCPro2 forms PD-localized punctate inclusions in virus-infected cells

To investigate the cellular compartment distribution of HCPro2 in virus-infected cells, we employed an ANRSV infectious cDNA clone – pRS as the backbone (Wang et al., 2021) to fuse a complete GFP-coding sequence at the beginning of HCPro2 and obtained the clone pRS-GFP-HCPro2 (Fig. 3A), followed by infectivity test in *N. benthamiana*. Eight days later, all inoculated plants (*n* = 10) exhibited chlorosis symptoms and obvious green fluorescence signals along veins in upper non-inoculated leaves (Fig. 3B), indicating that the recombinant virus clone is aggressive in *N. benthamiana*. In line, the fused protein GFP-HCPro2 (61.21 kDa) was detected (Fig. 3C). Subsequently, virus-infected leaf tissues were sampled for subcellular fractionation assay. Immunoblot analysis revealed that GFP-HCPro2 was present in different fractions with a varied degree, including nuclei-chloroplast-cell wall fraction (P3), membranous fraction (P30) and cytoplasmic fraction (S30) (Fig. 3D). As a control, the free GFP produced in pRS-G sample mainly existed in S30 fraction (Fig. 3D).

**Fig. 3.**
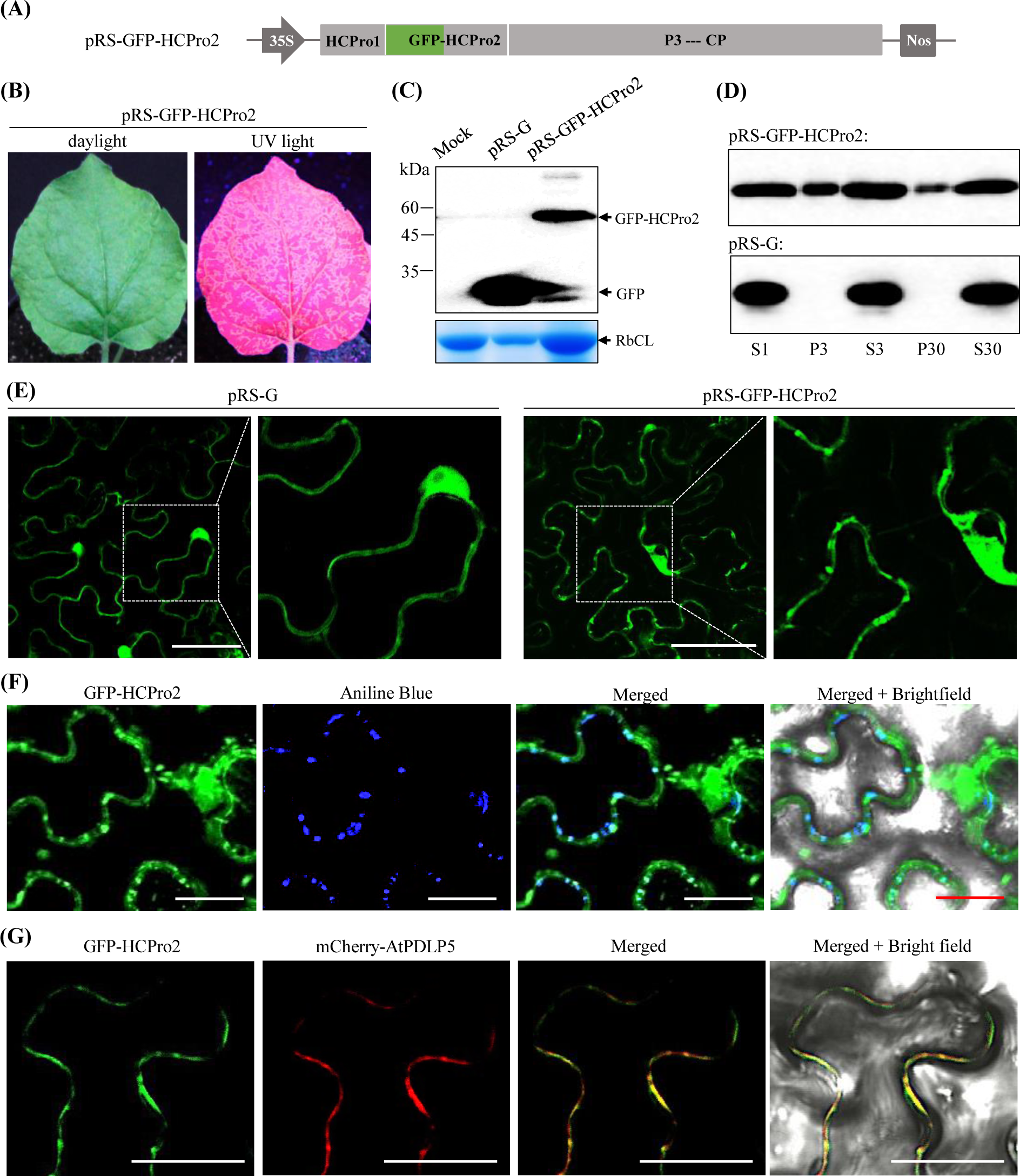
Cellular compartment distribution and subcellular localization of HCPro2 in virus-infected cells. **(A)** Schematic diagram of pRS-GFP-HCPro2. For the clone, the complete GFP-coding sequence was fused at the beginning of HCPro2. **(B)** Infectivity test of pRS-GFP-HCPro2 in *N. benthamiana*. The upper non-inoculated leaf was photographed under either daylight or UV light at 8 dpi. **(C)** Immunoblot detection of GFP-HCPro2 accumulation. The upper non-inoculated leaves of *N. benthamiana* plants infiltrated with either pRS-GFP-HCPro2 or pRS-G were assayed by Western blot at 8 dpi. Coomassie blue staining of RbCL was used as a loading control. **(D)** Subcellular fractionation coupled with immunoblot detection of GFP-HCPro2. The upper non-inoculated leaves of *N. benthamiana* plants infiltrated with either pRS-GFP-HCPro2 or pRS-G were collected at 8 dpi for subcellular fractionation assay. The resulting fractions were subjected to immunoblot detection of GFP-HCPro2 and free GFP. S1, the supernatant following centrifugation of crude homogenate at 1000 *g*; S3 and P3, the corresponding supernatant and pellet following the centrifugation of S1 at 3700 *g*; S30 and P30, the corresponding supernatant and pellet following the centrifugation of S3 at 30000 *g*. **(E)** Subcellular localization of GFP-HCPro2 in virus-infected cells. *N. benthamiana* leaves were inoculated with either pRS-GFP-HCPro2 or pRS-G, followed by confocal microscopy observation at 72 hpi. The regions indicated by dashed boxes are enlarged. Bars, 50 μm. **(F)** Subcellular co-localization of GFP-HCPro2 and the callose at PD. At 72 hpi, the inoculated leaves with pRS-GFP-HCPro2 were stained with aniline blue, followed by confocal microscopy observation. Bars, 25 μm. **(G)** Subcellular co-localization of GFP-HCPro2 and mCherry-AtPDLP5 (a PD marker). *N. benthamiana* leaves were co-inoculated with pCaM-GFP-HCPro2 and a plasmid for expressing mCherry-AtPDLP5, followed by confocal microscopy observation at 72 hpi. Bars, 25 μm.

Next, we examined the subcellular localization pattern of HCPro2 in virus-infected cells. *N. benthamiana* leaves infiltrated with either pRS-GFP-HCPro2 or pRS-G (OD_600_ = 0.1) at 72 hpi were subjected to confocal microscopy observation. A clear difference was observed between pRS-G and pRS-GFP-HCPro2 samples. Similar with the distribution pattern of free GFP in pRS-G sample, GFP-HCPro2 is diffused into cytoplasm and nucleus (Fig. 3E). However, GFP-HCPro2 accumulated in punctate structures, and the distribution pattern assembles that of PD-localized markers (Fig. 3E; Wei et al., 2010). To test this idea, leaf samples of pRS-GFP-HCPro2 were stained with aniline blue, which reacts with the callose deposited at the necks of PD. As expected, about 70% of GFP-HCPro2 inclusions co-localized with aniline blue-stained callose (Fig. 3F). In addition, expression of GFP-HCPro2 alone forms punctate inclusions as well, which co-localized with a PD maker - mCherry-AtPDLP5 (Fig. 3G).

In conclusion, the distribution of HCPro2 into different cellular compartments suggests that it might implement multiple functions in viral infection cycle. In particular, HCPro2 forms PD-localized inclusions, providing an important clue of the involvement of HCPro2 in viral cell-to-cell movement.

### Purification and identification of viral and host proteins that physically associate with HCPro2 in the context of viral infection

To get insight into the role of HCPro2 in viral intercellular movement, a twin-Strep sequence (2×Strep) was fused with the first nucleotide of HCPro2 in pRS-G (Fig. 4A) to purify and identity viral and host proteins that physically associate with HCPro2 in the context of viral infection. Infectivity test showed that pRS-G-2×Strep-HCPro2 is viable, but much weaker than the wild-type pRS-G (Fig. 4B). The fused 2×Strep-HCPro2 (37.57 kDa) was detected from upper non-inoculated leaves (Fig. 4C). Unfortunately, affinity purification with streptavidin failed to enrich 2×Strep-HCPro2 together with its associating proteins via sodium dodecyl-sulfate polyacrylamide gel electrophoresis (SDS-PAGE) analysis and immunoblot detection. Considered that the deletion of HCPro1 significantly increases either viral RNA load or protein expression (Fig. 1), we used pRS-G(ΔHCPro1) as the backbone instead to fuse twin-Strep with HCPro2 and obtained pRS-G(ΔHCPro1)-2×Strep-HCPro2 (Fig. 4A). Infectivity test showed that the clone was more aggressive than pRS-G and pRS-G-2×Strep-HCPro2 in either virus-triggered symptom phenotype or systemic spreading (Fig. 4B). A significant higher abundance of 2×Strep-HCPro2 was detected (Fig. 4C). Subsequently, the upper non-inoculated leaves of plants infiltrated with pRS-G(ΔHCPro1)-2×Strep-HCPro2 or pRS-G (as the parallel control) were subjected to affinity purification. SDS-PAGE analysis revealed the presence of a putative band corresponding to 2×Strep-HCPro2 and several other bands in co-purified products, and these bands were absent in the parallel control (Fig. 4D). An obvious band representing 2×Strep-HCPro2 was immuno-detected (Fig. 4D). Intriguingly, another band likely representing a 2×Strep-HCPro2-containing complex with a high-molecular-mess was immuno-detected as well (Fig. 4D). Liquid chromatography tandem mass spectrometry (LC-MS/MS) analysis identified six viral proteins (HCPro2, P3, 6K1, CI, NIb and CP) and a total of 63 host proteins in co-purified products, with the bait HCPro2 as the most abundant one (Fig. 4E, F; Supplemental Table S1).

**Fig. 4.**
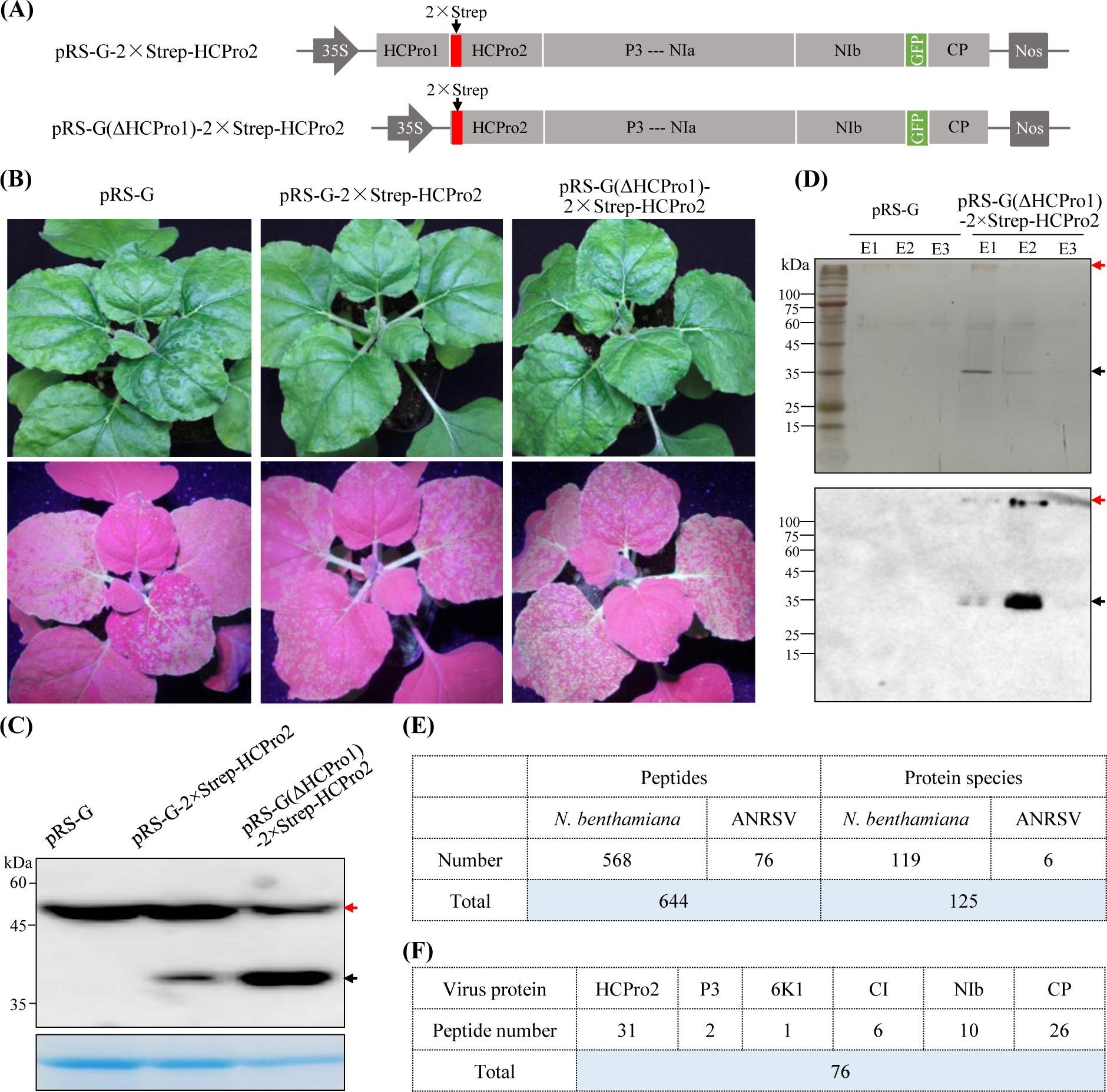
Purification and identification of viral and host proteins that associate with HCPro2 during ANRSV infection. **(A)** Schematic diagrams of pRS-G-2×Strep-HCPro2 and pRS-G(△HCPro1)-2×Strep-HCPro2. **(B)** Infectivity test of the indicated clones in *N. benthamiana*. The representative plants were photographed at 12 dpi. **(C)** Immunoblot detection of 2×Strep-HCPro2 in upper non-inoculated leaves at 12 dpi. Coomassie blue staining of RbCL was used as a loading control. **(D)** SDS-PAGE analysis and immunoblot detection of co-purified proteins with 2×Strep-HCPro2 as the bait. The upper non-inoculated leaves of plants infiltrated with either pRS-G (as the parallel control) or pRS-G(△HCPro1)-2×Strep-HCPro2 were collected at 12 dpi for affinity-purification with streptavidin. Elution fractions (E1-E3) were used for SDS-PAGE with silver staining (upper panel) and immunoblot detection (lower). The black arrows indicated a putative band corresponding to 2×Strep-HCPro2 (37.57 kDa). The bands (indicated by red arrows) likely represent a HCPro2-containing complex with high-molecular-mess. **(E)** Summary of the peptides and corresponding proteins. The peptides were analyzed by LC-MS/MS from co-purified products with 2×Strep-HCPro2. **(F)** Summary of viral proteins co-purified with 2×Strep-HCPro2.

### HCPro2 interacts with CI and CP *in planta*

Both CI and CP, known to be viral movement-related proteins for potyvirids, are co-purified with HCPro2 (Fig. 4F), promoting us to speculate that HCPro2 might regulate viral intercellular movement via interactions with CI and CP. To test this idea, we examined the interactions of HCPro2 with three viral movement-related factors (CI, CP and P3N-PIPO) using yeast two hybrid (Y2H). The coding sequences of them were respectively cloned into pGBKT7-DEST or pGADT7-DEST for the expression of GAL4 DNA binding domain (BD)- or activation domain (AD)-fused viral proteins. Co-transformation of yeast cells did not detect the interaction between BD-HCPro2 and each of AD-CI, AD-CP and AD-P3N-PIPO. Similarly, no interaction was observed when co-expressing AD-HCPro2 and each of BD-CI, BD-CP and BD-P3N-PIPO (Fig. 5A). Also, HCPro2 did not interact with the remaining viral factors in Y2H (Supplemental Fig. S3).

**Fig. 5.**
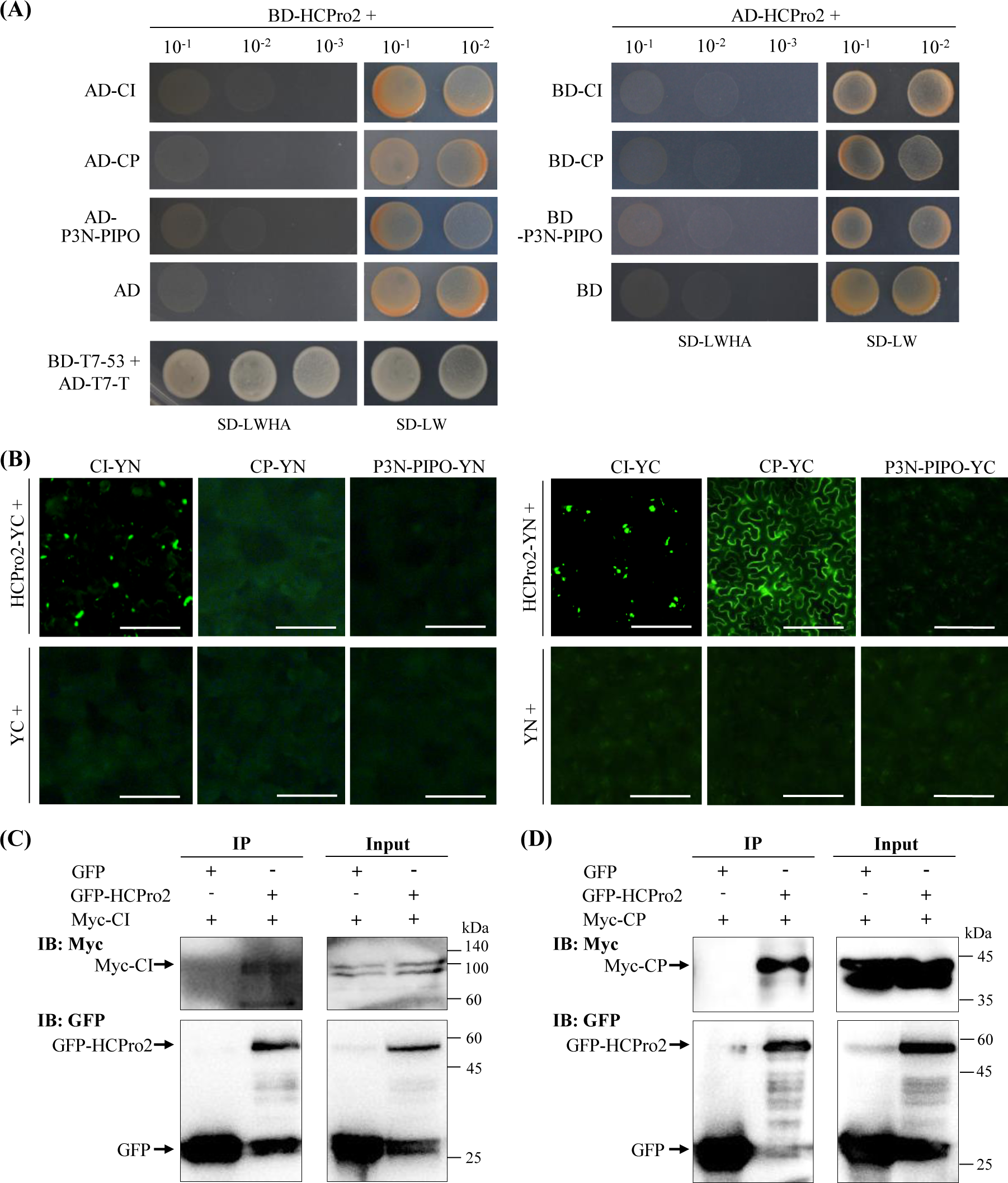
HCPro2 interacts with CI and CP *in vivo*. **(A)** Y2H assay test that interaction of HCPro2 with P3N-PIPO, CI or CP. The coding sequences of HCPro2, P3N-PIPO, CI and CP were respectively cloned into pGBKT7-DEST or pGADT7-DEST for the expression of these proteins fused with GAL4 BD or AD domain. Yeast competent cells (Y2H Gold) were co-transformed to express the indicated pairs of proteins. After incubation, the transformed cells were subjected to 10-fold serial dilutions and plated on the SD/-Trp/-Leu and SD/-Trp/-Leu/-His/-Ade mediums. The plates were cultured at 28 ℃ for four to six days before photographing. Co-transformation of a pair of plasmids for the expression of AD-T7-T and BD-T7-53 was included as the positive control. **(B)** The interactions of HCPro2 with CI, CP and P3N-PIPO tested by BiFC. The corresponding coding sequences of HCPro2, CI, CP and P3N-PIPO were engineered into p35S-gatewayYN or p35S-gatewayYC for the expression of these proteins fused with the YN or YC part of YFP. *N. benthamiana* leaves were co-inoculated for the expression of the indicated pairs of proteins. YFP signals (shown in green) were observed by fluorescence microscope at 72 hpi. Bars, 50 μm. **(C, D)** The interactions of HCPro2 with CI and CP tested by Co-IP. The inoculated leaves of *N. benthamiana* plants for the co-expression of GFP-HCPro2 / Myc-CI **(C)** or GFP-HCPro2 / Myc-CP **(D)** were sampled at 72 hpi for Co-IP assay using GFP-Trap Agarose. Total protein extracts prior to (Input) and after immunoprecipitation (IP) were analyzed by immunoblotting using anti-Myc and anti-GFP antibodies.

Next, we examined whether HCPro2 interacts with CI, CP and P3N-PIPO *in planta* using bimolecular fluorescence complementation (BiFC). Their coding sequences were each engineered into p35S-gatewayYN or p35S-gatewayYC to express viral proteins fused with N-terminal half (YN) or C-terminal half of YFP (YC). HCPro2-YC was transiently co-expressed with each of P3N-PIPO-YN, CI-YN and CP-YN in *N. benthamiana* leaves. The fluorescence signals with punctate distribution were observed for co-expression of HCPro2-YC and CI-YN (Fig. 5B), indicating that HCPro2 interacts with CI *in planta*. A consistent result was obtained when using a combination of HCPro2-YN and CI-YC for the test (Fig. 5B). In addition, we observed strong fluorescence signals (diffused intracellularly) from leaf sample co-expressing HCPro2-YN and CP-YC (Fig. 5B). However, fluorescence signals were not discriminated from the samples co-expressing either HCPro2-YC / P3N-PIPO-YN or HCPro2-YN / P3N-PIPO-YC (Fig. 5B). As a result, HCPro2 interacts with CI and CP *in planta*.

We further tested the interactions of HCPro2 with CI and CP by coimmunoprecipitation (Co-IP). For this, we developed a series of T-DNA constructs for transient expression of free GFP, GFP-tagged HCPro2 (GFP-HCPro2), 4×Myc-tagged CI (Myc-CI) or 4×Myc-tagged CP (Myc-CP). GFP-HCPro2 was co-expressed with either Myc-CI or Myc-CP in *N. benthamiana* leaves. Co-expression of GFP and Myc-CI or Myc-CP was included as the parallel controls. Total proteins were extracted at 72 hpi, and co-immunoprecipitated with GFP-Trap Agarose, followed by immunoblot analysis. The results showed that either CI or CP was co-immunoprecipitated with GFP-HCPro2, but not with GFP in the parallel controls (Fig. 5C, D).

### HCPro2, CI and CP form the interactive complex at PD in viral infection

During potyvirid infection, CI forms conical structures at PD, and binds CP or virion binds to aid viral intercellular movement. HCPro2 forms PD-localized punctate inclusions in virus-infected cells (Fig. 3). Thus, we speculated that the interactions of HCPro2 with CI and CP occur at PD in viral infection. To test this hypothesis, two constructs for the co-expression of HCPro2-YN and CI-YC, along with ANRSV infectious cDNA clone (pRS), were co-inoculated into *N. benthamiana* leaves. At 72 hpi, obvious punctate structures emitting yellow fluorescence, an indication of HCPro2 and CI interaction, were observed to overlap with aniline blue-strained callose at PD (Fig. 6A). Similarly, HCPro2-YN and CP-YC interact to form punctate structures either, which co-localized with the callose at PD (Fig. 6B). Next, the co-localization of HCPro2 and either CI or CP was examined in the presence of viral infection. For this, we produced two T-DNA constructs for the respective expression of mCherry-fused CI (CI-mCherry) and CP (CP-mCherry). Each of them, along with pRS-GFP-HCPro2, were co-inoculated into *N. benthamiana* leaves. The punctate structures of GFP-HCPro2 (resulting from viral genome expression), aniline blue-strained callose, and either CI-mCherry or CP-mCherry inclusions were largely overlapped at 72 hpi (Supplemental Fig. S4). Taken together, HCPro2 interacts with and colocalizes with CI and CP at PD in the presence of viral infection.

**Fig. 6.**
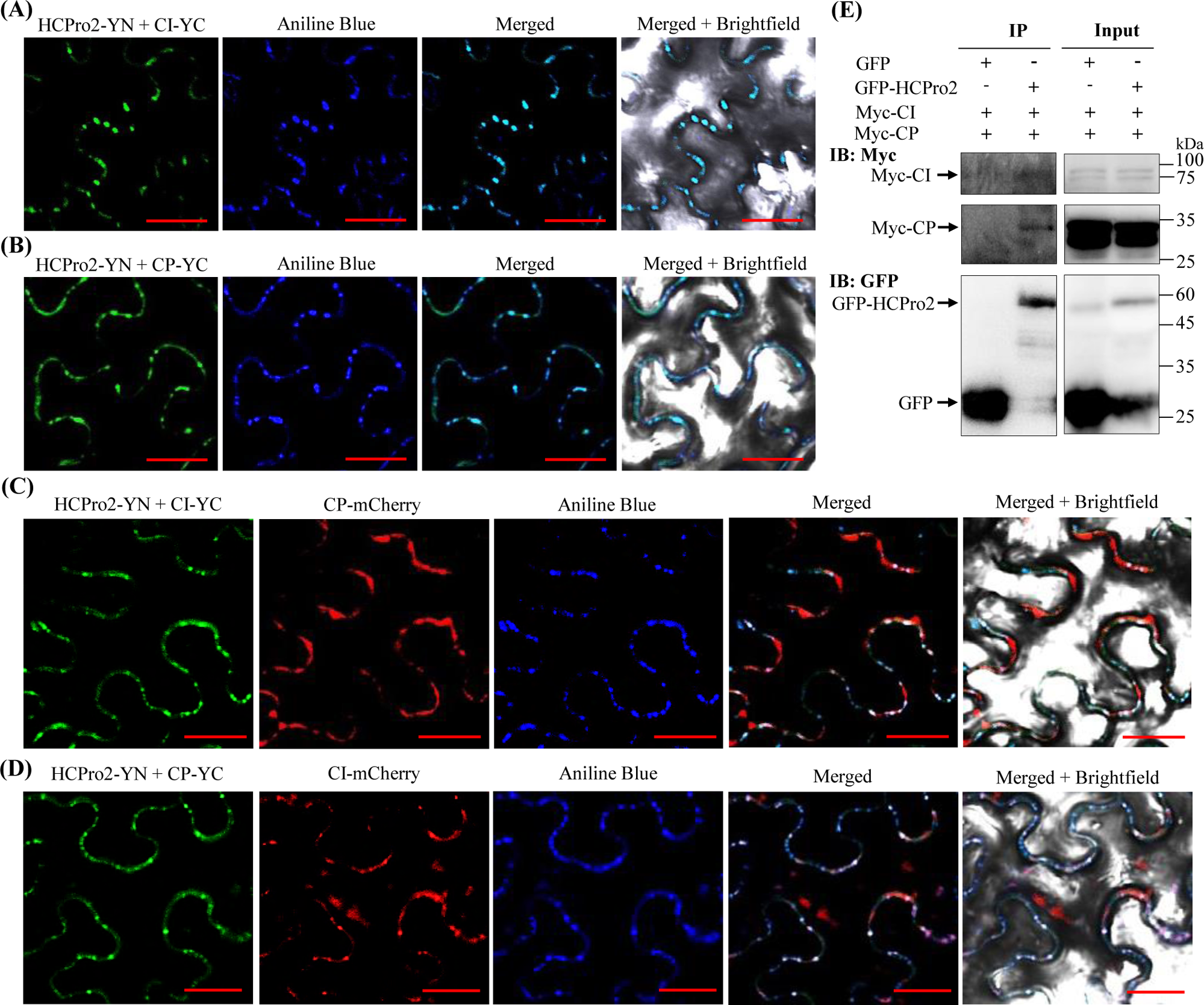
HCPro2-CI-CP forms the interactive complex at PD in viral infection. **(A)** HCPro2 interacts with CI at PD. *N*. *benthamiana* leaves were co-inoculated with two constructs corresponding to HCPro2-YN and CI-YC together with the viral clone – pRS (final OD_600_ = 0.2 per clone), followed by staining with aniline blue at 72 hpi and immediate observation by confocal microscopy. Bars, 25 μm. **(B)** HCPro2 interacts with CP at PD. *N. benthamiana* leaves were co-inoculated with two constructs corresponding to HCPro2-YN and CP-YC together with pRS (final OD_600_ = 0.2 per clone), followed by staining with aniline blue at 72 hpi and immediate observation by confocal microscopy. Bars, 25 μm. **(C)** Confocal microscopy observation of *N. benthamiana* leaves co-expressing HCPro2-YN, CI-YC and CP-mCherry in viral infection. *N. benthamiana* leaves were co-inoculated with three constructs for simultaneous expression of HCPro2-YN, CI-YC, and CP-mCherry together with pRS (final OD_600_ = 0.2 per clone), followed by staining with aniline blue at 72 hpi and immediate observation by confocal microscopy. Bars, 25 μm. **(D)** Confocal microscopy observation of *N. benthamiana* leaves co-expressing HCPro2-YN, CP-YC and CI-mCherry in viral infection. *N. benthamiana* leaves were co-inoculated with three constructs for simultaneous expression of HCPro2-YN, CP-YC and CI-mCherry along with pRS (final OD_600_ = 0.2 per clone), followed by staining with aniline blue at 72 hpi and immediate observation by confocal microscopy. Bars, 25 μm. **(E)** Both CI and CP were coimmunoprecipitated with GFP-HCPro2 in viral infection. *N. benthamiana* leaves are co-inoculated with two constructs for simultaneous expression of Myc-CI and Myc-CP along with the viral clone pRS-GFP-HCPro2. At 72 hpi, total proteins were extracted for a Co-IP assay using GFP-Trap Agarose. Total protein extracts prior to (Input) and after immunoprecipitation (IP) were immuno-detected using anti-Myc and anti-GFP polyclonal antibodies.

Further, we investigated whether the three viral factors (HCPro2, CI and CP) form interactive complex at PD in viral infection. Three constructs for co-expression of HCPro2-YN, CI-YC, and CP-mCherry, together with viral clone - pRS, were co-inoculated into *N. benthamiana* leaves. At 72 hpi, the punctate structures emitting green fluorescence, an indication of HCPro2 and CI interaction, overlapped with the structures formed by CP-mCherry at PD, albeit not completely (Fig. 6C). A similar result was obtained when co-expressing HCPro2-YN, CP-YC and CI-mCherry (Fig. 6D). The above results indicate that HCPro2-CI-CP forms the interactive complex at PD in viral infection. To further reveal the existence of HCPro2-CI-CP complex, a Co-IP assay was performed. Two constructs corresponding to Myc-CI and Myc-CP, together with pRS-GFP-HCPro2 or pRS-G (negative control) were co-inoculated into *N. benthamiana* leaves. Total proteins were extracted at 96 hpi, followed by immunoprecipitation using GFP-Trap Agarose. Immunoblot analysis showed that both CI and CP were coimmunoprecipitated by GFP-HCPro2 (Fig. 6E). In conclusion, the three movement-related proteins (HCPro2, CI and CP) interact with each other to form the complexes at PD in viral infection.

### A common host protein (NbRbCS) mediates the interactions of HCPro2-CI, HCPro2-CP, and CI-CP

Given that HCPro2 interacts with CI and CP *in planta* rather than *in vitro*, we proposed that one or more unknown host proteins mediate these interactions. To test this hypothesis, Y2H assay was employed to screen the interactions between HCPro2 and its associating host proteins. The candidate proteins with the score above 25 by LC-MS/MS (Supplemental Table S1) were selected. Y2H assays revealed a strong interaction between HCPro2 and NbRbCS (Fig. 7A). The interaction was further confirmed by BiFC and Co-IP (Fig. 7B, C). Intriguingly, a high-molecular-mess protein likely representing GFP-HCPro2- and NbRbCS-containing complex was detected in co-immunoprecipitated products by GFP-HCPro2 (Fig. 7C). A similar phenomenon was observed when performing affinity-purification with 2×Strep-HCPro2 as the bait in viral infection (Fig. 4D). These observations suggest that an interactive complex, at least including HCPro2 and NbRbCS, is prone to being formed *in planta*. Further, it was found that either N-terminal region (N2) or C-terminal cysteine protease region (D2) of HCPro2 interacts with NbRbCS (Supplemental Fig. S5).

**Fig. 7.**
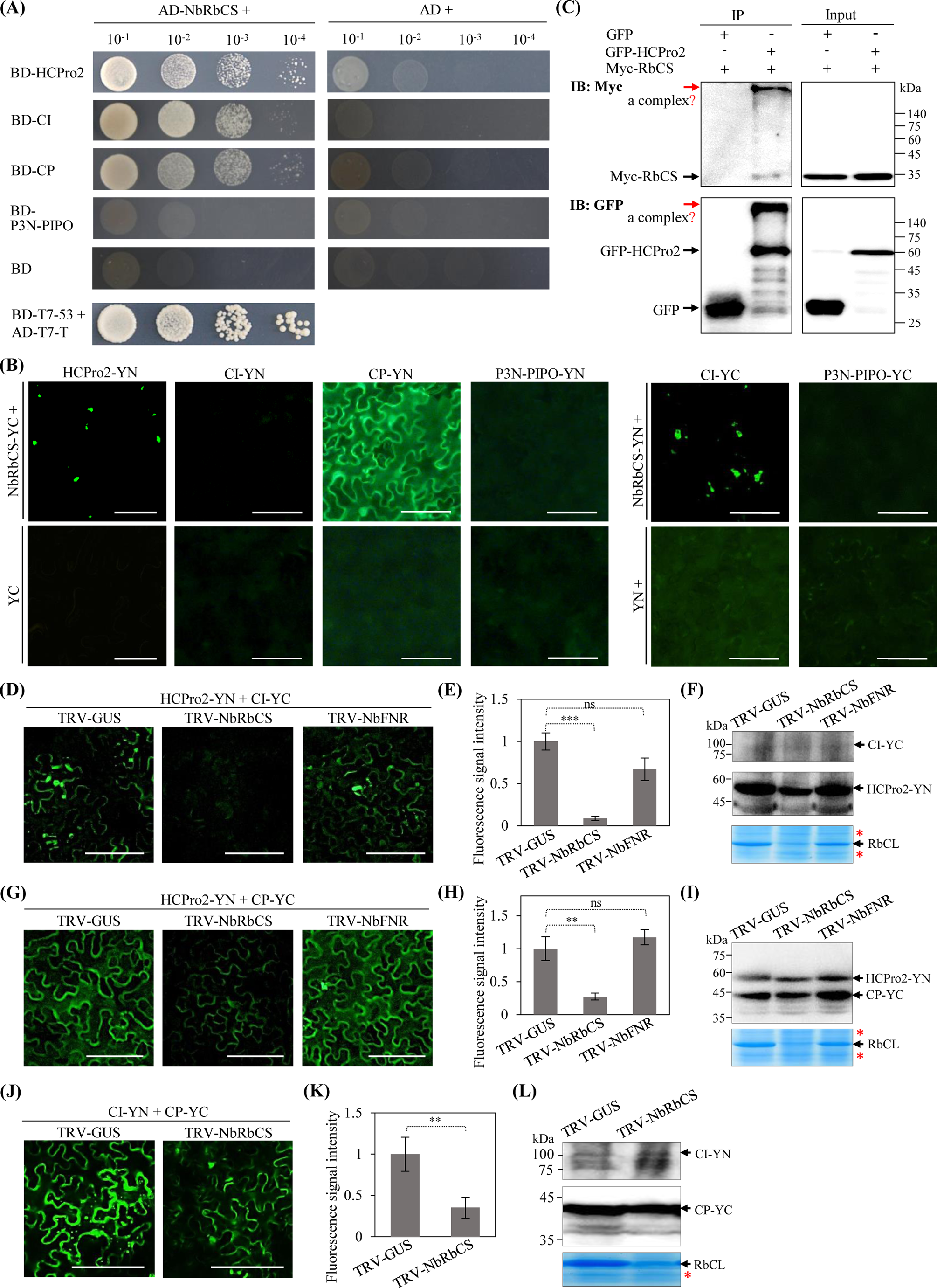
A common host protein (NbRbCS) mediates the interactions of HCPro2-CI, HCPro2-CP and CI-CP. **(A)** Y2H tests the interactions of NbRbCS with HCPro2, CI, CP and P3N-PIPO. The coding sequence of NbRbCS was cloned into pGADT7-DEST for expressing GAL4 AD-fused NbRbCS (AD-NbRbCS). Yeast cells were co-transformed to express a pair of indicated proteins, followed by 10-fold serial dilutions and plating on SD/-Trp/-Leu/-His/-Ade medium. The plates were placed at 28 ℃ for four to six days before photographing. Co-transformation of a pair of plasmids for the expression of AD-T7-T and BD-T7-53 was included as the positive control. **(B)** BiFC tests the interactions of NbRbCS with HCPro2, CI, CP, and P3N-PIPO. The coding sequence of NbRbCS was integrated into p35S-gatewayYN or p35S-gatewayYC for expressing YFP YN- or YC-fused NbRbCS (NbRbCS-YN or NbRbCS-YC). *N. benthamiana* leaves were co-inoculated for expressing a pair of indicated proteins (final OD_600_ = 0.2 per plasmid). The fluorescence signals (indicated in green color) were observed by a fluorescence microscope at 72 hpi. Bars, 50 μm. The co-expression of YC or YN along with an indicated protein was included as the negative control. **(C)** Co-IP tests the interaction between NbRbCS and HCPro2. *N. benthamiana* leaves for co-expression of Myc-NbRbCS and GFP-HCPro2 or GFP (final OD_600_ = 0.3 per plasmid) were sampled at 72 hpi for Co-IP assay using GFP-Trap Agarose. Total protein extracts prior to (Input) and after immunoprecipitation (IP) were analyzed by immunoblotting with anti-Myc and anti-GFP polyclonal antibodies. The bands indicated by red arrows represent a putative complex that at least contains GFP-HCPro2 and Myc-NbRbCS. **(D, G, J)** BiFC assays test the interactions of HCPro2-CI, HCPro2-CP, and CI-CP in *NbRbCS*- or *NbFNR*-silenced *N. benthamiana* plants. pTRV1, along with pTRV2-GUS (TRV-GUS), pTRV2-NbRbCS (TRV-NbRbCS) or pTRV2-NbFNR (TRV-NbFNR) were co-inoculated into *N. benthamiana* seedlings. At 12 dpi, the upper fully-expanded leaves were co-inoculated for co-expression of HCPro2-YN / CI-YC **(D)**, HCPro2-YN / CP-YC **(G)**, or CI-YN / CP-YC **(J)**. The OD_600_ value for each plasmid is finally adjusted to 0.2. The samples were observed by confocal microscopy at 60 hpi (G) or 72 hpi (D, J). Bars, 100 μm (D, G) or 50 μm (J). **(E, H, K)** Statistical analysis of fluorescence signal intensity. The fluorescence signal intensity, reflecting the interactive degree for HCPro2-YN / CI-YC **(E)**, HCPro2-YN / CP-YC **(H)**, or CI-YN / CP-YC **(K)**, was quantified by ImageJ. At least 20 scans per treatment from three independent experiments were analyzed. Data are presented as the mean ± SD (*n* ≥ 20). ***, *P*<0.001; **, 0.001<*P*<0.01; ns, no significant difference. **(F, I, L)** Immunoblot analysis of the accumulation of indicated proteins in co-inoculated leaves. The co-inoculated leaves for co-expression of HCPro2-YN / CI-YC **(F)**, HCPro2-YN / CP-YC **(I)** or CI-YN / CP-YC **(L)** were sampled at 60 hpi (I) or 72 hpi (F, L) for immunoblot analysis using anti-GFP antibody. As the abundance of RbCL was greatly decreased along with *RbCS*-silencing, the coomassie blue staining of protein bands (indicated by red asterisks) was used as a loading control.

Subsequently, we tested whether NbRbCS interacts with CI or CP. Y2H assays showed that AD-NbRbCS interacts with either BD-CI or BD-CP (Fig. 7A). The interactions were verified by BiFC (Fig. 7B). Moreover, NbRbCS was tested to not interact with P3N-PIPO by either Y2H or BiFC (Fig. 7A, B). In addition, we performed Y2H assays to examine the interactions of NbRbCS with the remaining viral factors (HCPro1, P3, 6K1, 6K2, VPg, NIa-Pro and NIb). Remarkably, the strong interactions of NbRbCS with HCPro1, P3, VPg and NIb were detected as well (Supplemental Fig. S6). Collectively, NbRbCS might function as a common host factor in mediating the interactions of HCPro2 with CI and CP.

To further illustrate the role of NbRbCS in mediating these interactions, we employed tobacco rattle virus (TRV)-based virus-induced gene silencing (VIGS) to knockdown *NbRbCS* expression in *N. benthamiana*. Compared with *N. benthamiana* plants inoculated with TRV-GUS (as a control), the plants inoculated with TRV-NbRbCS exhibited abnormal development phenotype such as dwarfism in size and foliar yellowing at 12 dpi (Supplemental Fig. S7). Real-time RT-qPCR confirmed that *NbRbCS* mRNA transcripts are significantly reduced in plants inoculated with TRV-NbRbCS (Supplemental Fig. S7). At this time point, upper non-inoculated leaves of *NbRbCS*-silenced and control plants were subjected to co-expression of HCPro2-YN and CI-YC. At 72 hpi, strong fluorescence signals, an indication of HCPro2 and CI interaction, were monitored for control plants, whereas the signals were nearly undetectable in *NbRbCS*-silenced plants (Fig. 7D, E). Either HCPro2-YN or CI-YC in *NbRbCS*-silenced plants accumulates at a comparable level with those in control plants (Fig. 7F). The abundance of RbCL is controlled by its interaction with RbCS to form L_8_S_8_ complex (Wietrzynski et al., 2021). Supporting this notion, we observed that NbRbCL is less accumulated in *NbRbCS*-silenced plants (lower panel in Fig. 7F). Similarly, the interaction between HCPro2 and CP was significantly attenuated in *NbRbCS*-silenced plants (Figure 7G-I). As known, silencing of *NbRbCS* destroys photosynthetic pathway, leading to abnormal physiological phenotype (Supplemental Fig. S7). To discriminate whether the effects of NbRbCS on HCPro2-CI and HCPro2-CP interactions were caused by the deficiency-of-photosynthesis, we silenced another key gene - Ferredoxin-NADP reductase gene (*FNR*) in photosynthetic pathway. As shown, silencing of *NbFNR* contributes to similar abnormalities with those observed in *NbRbCS*-silenced plants (Supplemental Fig. S7A, C). BiFC assays showed that silencing of *NbFNR* did not affect the interactions of HCPro2 with either CI or CP, which was in contrast to what was observed in *NbRbCS*-silenced plants (Fig. 7D-I).

For potyvirids, CP or virion binds with CI-forming conical structures at PD to facilitate viral cell-to-cell movement (Rodriguez-Cerezo et al., 1997; Roberts et al., 1998; Gabrenaite-Verkhovskaya et al., 2008). However, the interaction between CI and CP was detected *in planta* rather than *in vitro* in most cases (Guo et al., 2001; López et al., 2001; Kang et al., 2004; Lin et al., 2009; Zilian & Maiss, 2011; Deng et al., 2015). Considered that NbRbCS interacts with either CI or CP (Fig. 7A, B), we envisaged that NbRbCS might mediate the interaction between CI and CP. To test this hypothesis, CI-YN and CP-YC were co-expressed in upper non-inoculated leaves of *NbRbCS*-silenced or control plants. At 72 hpi, strong fluorescence signals, an indication of CI and CP interaction, were observed in control samples (Figure 7J, K). However, this interaction was significantly compromised in *NbRbCS*-silenced plants (Figure 7J, K). Either CI-YN or CP-YC in *NbRbCS*-silenced plants accumulates at a comparable level with those in control plants (Fig. 7L). Altogether, NbRbCS acts as a common host protein to mediate the interactions of HCPro2-CI, HCPro2-CP, and CI-CP.

### Interactions of NbRbCS with HCPro2, CI and CP occur at PD in viral infection

HCPro2-CI-CP forms interactive complex at PD in viral infection, and the interactions among them are mediated by a common protein – NbRbCS (Figs 6, 7). These promoted us to speculate that NbRbCS interacts with the three viral factors at PD in viral infection. A pair of T-DNA constructs for the co-expression of RbCS and each of these viral factors, together with viral clone – pRS, were co-inoculated into *N. benthamiana* leaves, followed by aniline blue staining at 72 hpi. Confocal microscopy revealed that the interactions of NbRbCS with HCPro2, CI or CP consistently punctate inclusions, which are substantially overlapped with the aniline blue-stained callose at PD (Fig. 8A-C). RbCS is a nucleus-encoded protein and transported into chloroplast via its N-terminal transit peptide (Grimm et al., 1997; Whitney et al., 2011). Hence, we examined whether NbRbCS was recruited to PD during viral infection. For this, we generated a construct for expressing a mCherry-tagged NbRbCS (NbRbCS-mCherry). As shown in Fig. 8D, NbRbCS-mCherry, when expressed alone, exactly localized at chloroplast, but not PD at all (Fig. 8D). Intriguingly, the co-localization of NbRbCS-mCherry with alanine blue-stained callose at PD was obviously observed, as long as NbRbCS-mCherry and viral clone - pRS were co-expressed in *N. benthamiana* leaves (Fig. 8E, F).

**Fig. 8.**
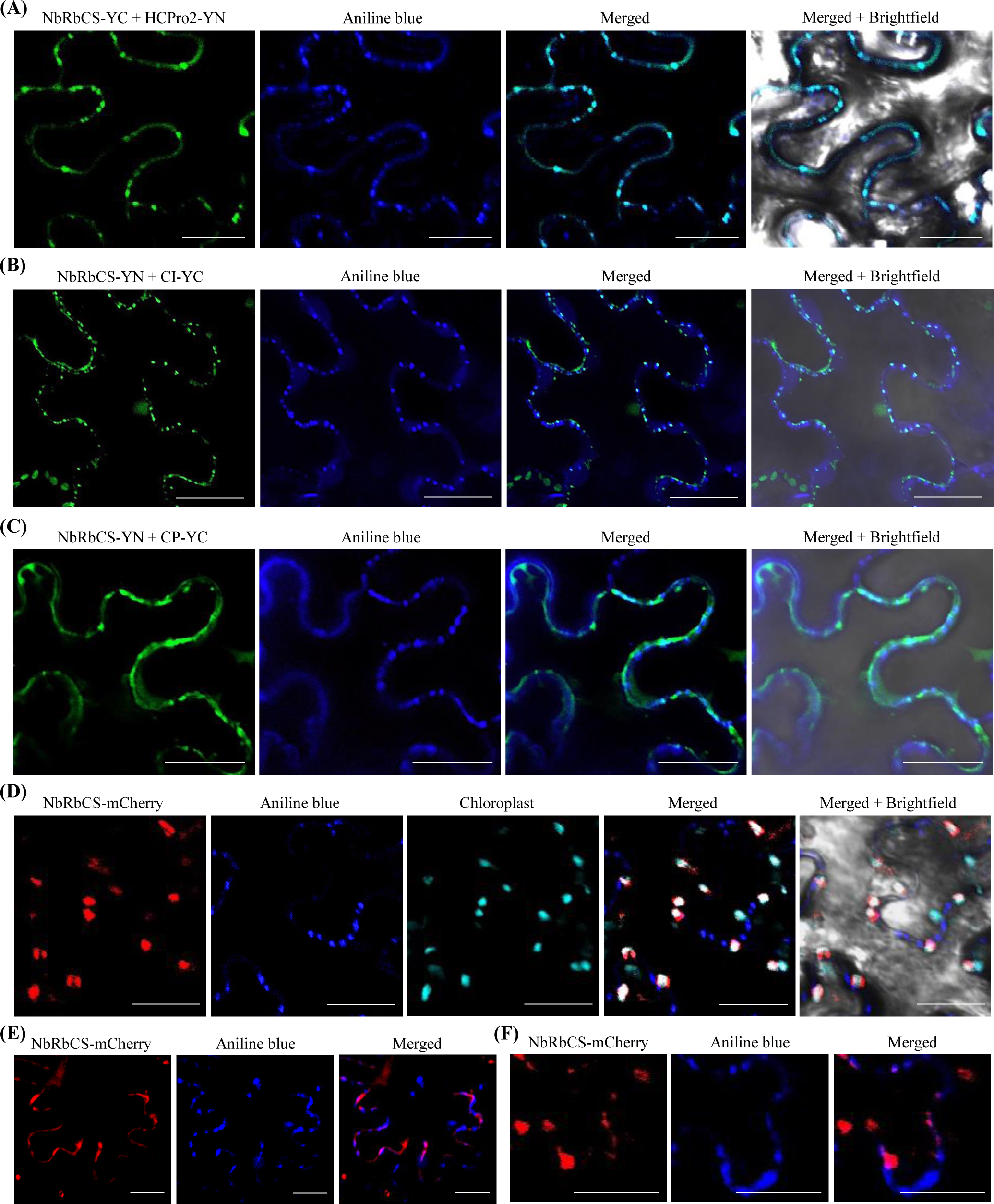
NbRbCS interacts with HCPro2, CI and CP at PD during viral infection. **(A-C)** Subcellular localization of interactive sites between NbRbCS and HCPro2, CI or CP in viral infection. A pair of constructs for co-expression of NbRbCS-YC / HCPro2-YN **(A)**, NbRbCS-YN / CI-YC **(B)**, or NbRbCS-YN / CP-YC **(C),** together with viral clone – pRS were co-inoculated into *N. benthamiana* leaves (final OD_600_ = 0.2 per clone). The inoculated leaves were stained with aniline blue at 72 hpi, immediately followed by confocal microscopy observation. Bars, 25 μm. **(D)** NbRbCS-mCherry targets chloroplast when expressed alone. *N. benthamiana* leaves were inoculated with the construct of NbRbCS-mCherry (OD_600_ = 0.2), followed by aniline blue staining at 72 hpi and confocal microscopy observation. Bars, 25 μm. **(E, F)** Subcellular localization of NbRbCS-mCherry at PD in viral infection. *N. benthamiana* leaves were co-expressed with NbRbCS-mCherry and viral clone (pRS) (OD_600_ = 0.2 per clone), followed by aniline blue staining at 72 hpi and confocal microscopy observation. A close-view of co-localization of NbRbCS-mCherry with aniline blue-stained callose at PD is shown in panel (F). Bars, 25 μm.

### Knockdown of *NbRbCS* significantly attenuates viral cell-to-cell movement and systemic infection for ANRSV and other three tested viruses in *Potyvirus* genus

Next, we investigated the effects of NbRbCS on ANRSV infection. *N. benthamiana* seedlings (*n* = 8 per clone) were pre-inoculated with TRV-NbRbCS, TRV-GUS or TRV-NbFNR (the parallel control). At 12 dpi, either *NbRbCS* or *NbFNR* mRNA transcripts were significantly reduced in corresponding plants (Supplemental Fig. S7). At this time point, these plants were challenged with ANRSV-GFP via sap rub-inoculation. Ten days later, strong green fluorescence signals, an indication of ANRSV-GFP infection, were observed in upper leaves of all plants pre-treated with TRV-GUS or TRV-NbFNR, whereas *NbRbCS*-silencing plants exhibited scattered fluorescence signals along veins in top leaves (Fig. 9A). Real-time RT-qPCR and Western blot assays confirmed that ANRSV infection was largely restricted in *NbRbCS*-silencing plants (Fig. 9B, C). We further examined the effects of *NbRbCS*-silencing on viral intercellular movement. For this, agrobacterial culture harboring pRS-G was highly diluted to OD_600_ of 0.001, and infiltrated into fully-expanded leaves of either control or *NbRbCS*-silenced plants. At 108 hpi, the size of viral infection foci resulting from viral spreading from primarily-transfected cells to peripheral cells was much smaller in *NbRbCS*-silenced leaves (Fig. 9D, E). Conclusively, knockdown of *NbRbCS* significantly debilitated viral infection, likely owing to restricting viral cell-to-cell movement.

**Fig. 9.**
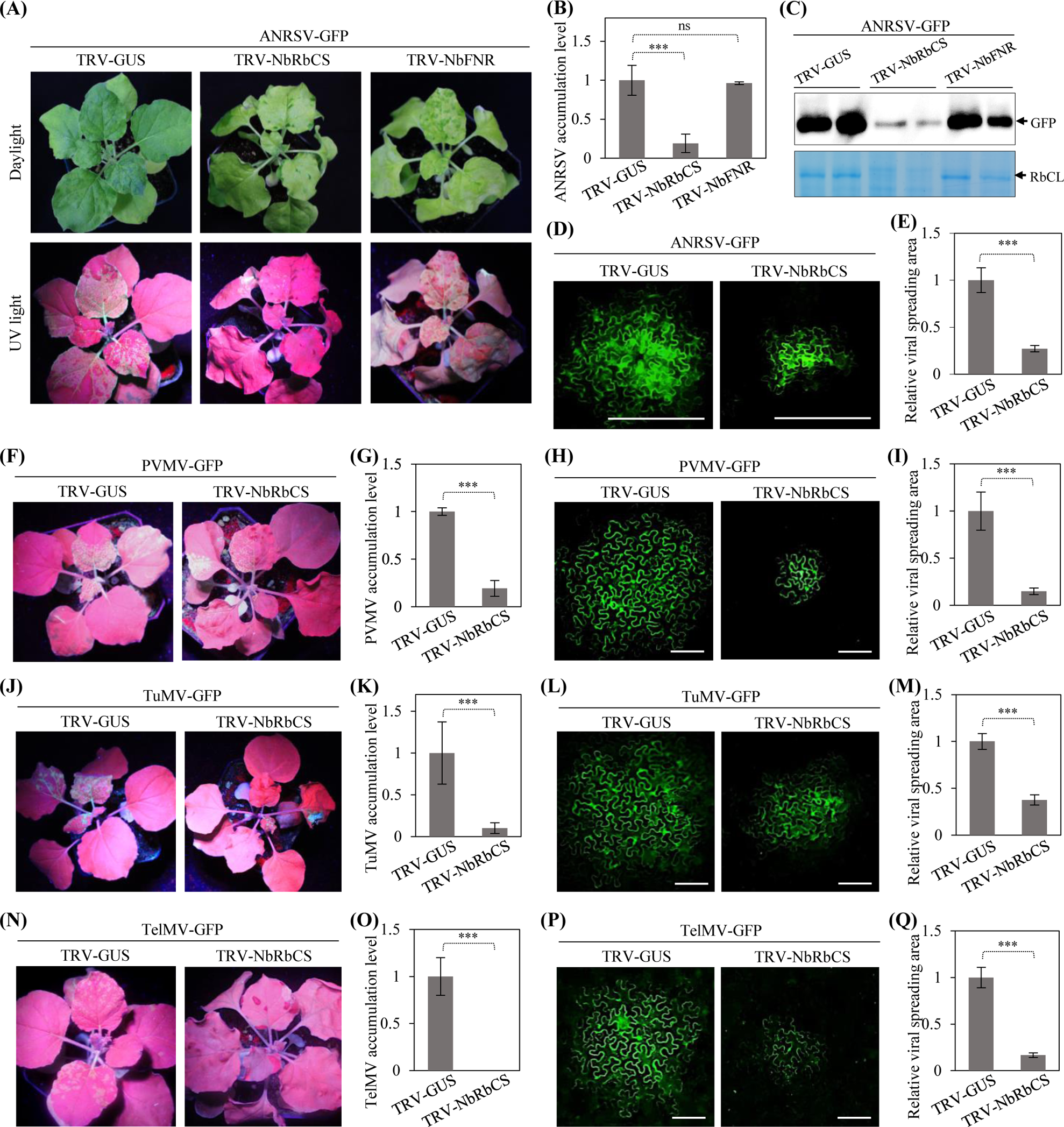
Knockdown of *NbRbCS* largely inhibits viral intercellular movement and systemic infection for ANRSV and other three tested potyviruses. (A) Silencing of *RbCS* significantly restricts ANRSV infection. *N. benthamiana* seedlings at 3- to 5-leaf stage were pre-inoculated with TRV-GUS, TRV-NbRbCS or TRV-NbFNR. At 12 dpi, these plants were challenged with ANRSV-GFP via sap sub inoculation. The representative plants were photographed under either daylight or UV light at ten days post-challenging inoculation (dpci). **(B)** Real-time RT-qPCR analysis of viral genomic RNA accumulation. Leaf samples were collected at 10 dpci for real-time RT-qPCR assay. Error bars denote the standard errors from three biological replicates. ***, *P*<0.001; ns, no significant difference. **(C)** Immunoblot analysis of GFP accumulation at 10 dpci. **(D)** Viral intercellular movement from single primarily infected cells at 108 hours post-challenging inoculation (hpci). Bars, 100 μm. **(E)** Statistical analysis of the size of viral infection foci at 108 hpci. A total of 25 infection foci per treatment from three independent experiments were analyzed by ImageJ. The size of infection foci is presented as the mean ± SD (*n* = 25). ***, *P*<0.001. **(F, K, N)** The effects of *RbCS*-silencing on the infectivity of three potyviruses. The representative plants were photographed under either daylight or UV light at 6 dpci for PVMV-GFP **(F)** and TuMV-GFP **(J)**, and at 13 dpci for TelMV-GFP **(N)**. **(G, K, O)** Real-time RT-qPCR analysis of viral genomic RNA accumulation. Viral genomic RNAs accumulation was determined at 6 dpci for PVMV-GFP **(G)** and TuMV-GFP **(K)**, and at 13 dpci for TelMV-GFP **(O)**. Error bars denote the standard errors from three biological replicates. ***, *P*<0.001. **(H, L, P)** Viral intercellular movement from single primarily infected cells. Viral intercellular movement was recorded at 108 hpci for PVMV-GFP **(H)** and at 84 hpci for TuMV-GFP **(L)** and TelMV-GFP **(P)**. Bars, 100 μm. **(I, M, Q)** Statistical analysis of the size of viral infection foci. The infection foci were determined at 108 hpci for PVMV-GFP **(I)** and at 84 hpci for TuMV-GFP **(M)** and TelMV-GFP **(Q).** A total of 20 infection foci per treatment from three independent experiments were analyzed. The size of infection foci is presented as the mean ± SD (*n* = 25). ***, *P*<0.001.

We employed a similar strategy to test the effects of NbRbCS on viral infectivity for other three viruses in *Potyvirus* genus, including pepper veinal mottle virus (PVMV), telosma mosaic virus (TelMV) and TuMV. The results showed that silencing of *NbRbCS* significantly weakened both systemic infection and cell-to-cell movement for PVMV (Fig. 9F-I), TuMV (Fig. 9J-M) and TelMV (Fig. 9N-Q). These data indicate that NbRbCS plays a general regulatory role in potyvirid infection.

## Discussion

Long time ago, it was observed that HCPro possesses the MP-related properties, but its connection with viral movement has been not demonstrated thus far. In this study, we provide genetic evidences in supporting the role of HCPro2 in intercellular movement, which could be functionally complemented by its counterpart from a potyvirus. The HCPro2, together with movement-related proteins CI and CP, form interactive complex at PD. The interactions among them are mediated by a common host protein – RbCS. Knockdown of *RbCS* greatly impairs these interactions, and viral cell-to-cell movement and systemic infection. Therefore, we envisage a scenario that RbCS, during its progressing to chloroplast, is hijacked as a pro-viral factor to mediate the assembly of intercellular movement complex to promote viral cell-to-cell movement (Fig. 10). The circumstance might be generally applied to other potyvirids, based on following considerations: i) The interactions among the three viral units (HCPro, CI and CP) have been documented for numerous potyvirids, but these interactions were usually detected *in planta*, but few detected *in vitro* (Langenberg, 1993; Rodriguez-Cerezo et al., 1997; Peng et al., 1998; Roberts et al., 1998; Guo et al., 1999; Guo et al., 2001; López et al., 2001; Roudet-Tavert et al., 2002; Kang et al., 2004; Gabrenaite-Verkhovskaya et al., 2008; Lin et al., 2009; Zilian & Maiss, 2011; Deng et al., 2015); ii) Silencing of *NbRbCS* significantly attenuates the cell-to-cell movement for either ANRSV or other three potyviruses tested.

**Fig. 10.**
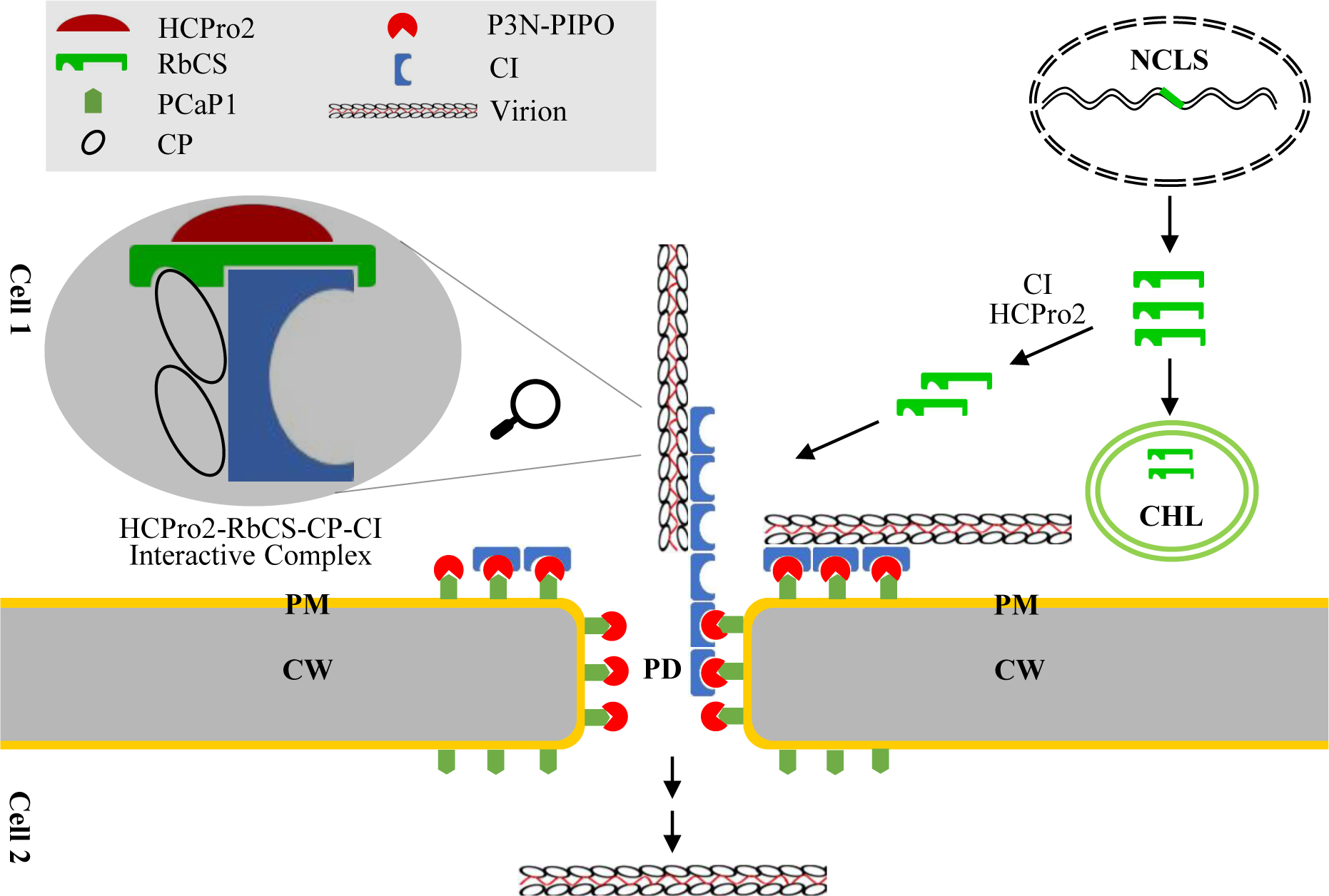
A working model reflecting that NbRbCS is co-opted as the scaffold protein in mediating the assembly of viral cell-to-cell movement complex. NCLS, nucleus; CHL, chloroplast; PM, plasma-membrane; CW, cell wall; PD, plasmodesmata.

The genomic 5′-terminal regions of potyvirids encode two types of leader proteases: serine-protease (P1) and cysteine-protease (HCPro), which differ greatly in the arrangement and sequence composition among inter-genus viruses (Cui & Wang, 2019; Yang et al., 2021; Pasin et al., 2022). For all tested potyvirids, one of the leader proteases expresses RSS activity. Two arepaviruses (ANRSV and ANSSV) have a distinguishably-arranged HCPro1-HCPro2. HCPro2 but not HCPro1 exerts the RSS function. HCPro1 is dispensable for ANRSV infection when tested in *N. benthamiana* (Fig. 1). The lethality of HCPro1 deletion in ANSSV (Qin et al., 2021) might be explained by the fact that *N. benthamiana* is much less susceptible to ANSSV infection (Wang et al., 2021). The phenomenon that the leader protease without RSS activity is dispensable for viral infectivity has been reported for several potyvirids (Stenger et al., 2005a; Pasin et al., 2014; You and Shirako, 2010). A clone of wheat streak mosaic virus (WSMV; *Tritimovirus*) with the deletion of entire HCPro sequence is still viable (Stenger et al., 2005a), whereas the HCPro with loss-of-RSS activity is a determinant in eriophyid mite-vectored transmission (Stenger et al., 2005b). In the case of plum pox virus (PPV; *Potyvirus*), the P1 protein is not essential for viral systemic infection, although it elaborately modulates viral replication to evade host immune response (Pasin et al., 2014). Here, we speculate that HCPro1, albeit not essential for viral infectivity, might play an accessory role in viral infection or function in virus-vectored transmission.

Potyvirid HCPro is likely involved in viral cell-to-cell movement. However, there is lacking of sufficient genetic and cytopathological evidences. This study comprehensively investigated its homolog in ANRSV (HCPro2), and provided following evidences to support its role in cell-to-cell movement: i) Replacement of HCPro2 with an unrelated VSR protein has no obvious effect on viral RNA accumulation, whereas nearly abolishes viral cell-to-cell movement (Fig.2). ii)Substitution of HCPro2 with its counterpart from a potyvirus efficiently complements viral intercellular movement, indicating that inter-genus HCPros are functionally interchangeable in aiding viral intercellular movement. ii) The movement-related proteins CI and CP are co-purified with HCPro2 in a context of viral infection. iii) HCPro2, CI and CP form PD-targeting interactive complex, which proves to be pivotal in viral cell-to-cell movement. The above results, together with previous observations (Rojas et al., 1997) and the fact that HCPro interacts with CI or/and CP *in planta* for numerous potyvirids (Valli et al., 2018; Revers & García, 2015; Sorel et al., 2014; Martínez-Turiño & García, 2020), suggest that HCPros among different potyvirids might share a common efficacy in aiding viral intercellular movement. Nevertheless, the underlying molecular mechanism is still unknown at this time. Previous studies revealed that HCPros of both PPV and PVA have the capacity to stabilize CP and enhance the yield of viral particles (Valli et al., 2014; Saha et al., 2023), suggesting that HCPro aids viral intercellular movement in an indirect manner. Intriguingly, the steady-state of CP mediated by HCPro was observed either in a context of viral infection or in the presence of viral proteins P3-to-CP (Valli et al., 2014; Saha et al., 2023), whereas the co-expression of HCPro and CP does not (Saha et al., 2023). Consequently, we propose that HCPro facilitates the stability of CP and viral particle yield via the formation of HCPro-RbCS-CP-CI interactive complex (Fig. 10).

HCPro2 is distributed, with a varied degree, into different cellular compartments in viral infection. The HCPro2-formed inclusions mainly target to PD, but a small portion of them are elsewhere. In recent years, a significant progress has been achieved with regard to the aggregates induced by PVA HCPro. The aggregates (called as PVA-induced granules, PGs) comprise of several host proteins and viral genomic RNAs, and are multifunctional during viral infection, including viral genome translation, RSS, encapsidation and systemic spread (Hafrén et al., 2015; De et al., 2020; Pollari et al., 2020). Whether the HCPro2 inclusions that are not targeted to PD behave similar functions with PGs await to be investigated. Among five viral proteins co-purified with HCPro2, four (P3, 6K1, CI and NIb) are components of 6K2-induced replication complex of potyvirids (Revers & García, 2015; Cui & Wang, 2019; Yang et al., 2021). As well, HCPro was identified from 6K2-induced replication vesicles in the case of PVA (Lõhmus et al., 2016). It is rational to speculate that HCPro2 might also participate in viral replication, which would be a promising research direction.

The chloroplast, an organelle only found in plant cells and some protists, has long been recognized as a common target by many plant viruses. Plant viruses may directly modify chloroplast membranes to assemble viral replication complex, or co-opt chloroplast proteins for viral replication, movement or/and counteracting host defense response. The rubisco is highly expressed in plants, and believed to be the most abundant protein on the planet (Bar-On & Milo, 2019). However, only one document is dedicated to the description of RbCS-virus interaction and its biological relevance (Zhao et al., 2013). In this study, we provide multi-discipline evidences to support the notion that NbRbCS is co-opted to mediate multiple interactions among viral movement-related proteins, likely functioning in the assembly of movement complex (Fig. 10). Here, we discuss three critical points that need to be clarified in future: i) How is chloroplast-localized RbCS recruited to PD? HCPro2, when expressed alone *in planta*, is able to target PD (Fig. 3G). Potyvirid CI is recruited by P3N-PIPO to form PD-localized cone-shaped structures (Wei et al., 2010). Thus, at least the two viral factor – CI and HCPro2 together recruit RbCS to PD in viral infection (Fig. 10). The CP or virions bind with CI cone structures of at PD, likely via the mediator – RbCS. ii) Whether the multiple interactions among HCPro2, RbCS, CP and CI (including potential self-interactions) have synergistic enhancement effect awaits to be investigated. iii) It is so fascinating that RbCS, such a small molecular, interacts with three viral movement-related proteins. In the RbCS-mediated complex, it is unclear whether one molecular of RbCS could simultaneously interact with HCPro2, CI and CP, or more molecular are needed. To clarify this point, a fine mapping of interaction sites between RbCS and HCPro2, CI or CP should be performed.

A previous report showed that RbCS interacts with P3 for several potyviruses (Lin et al., 2011). Besides P3, other three protein (HCPro1, VPg and NIb) of ANRSV interact with RbCS as well (Supplemental Fig. S6). Coincidentally, two of them (P3 and NIb) are co-purified with HCPro2. VPg plays multifunctional roles during viral infection, and when it is part of 6K2–VPg–NIa-Pro product, VPg is targeted to membranous factories induced by the virus where it plays a key role in viral RNA replication (Beauchemin et al., 2007; Wei & Wang, 2008). Taken together, we envisage that RbCS might also participate in viral replication via its interactions with replication-related viral proteins. Again, it is amazing that a redundant chloroplast protein has the capacity of interaction with multiple viral proteins. A fine mapping of interaction sites among them might facilitate to precisely design a broad-spectrum potyvirid-resistance strategy.

## Materials and methods

### Plant materials and virus resources

*N. benthamiana* plants used in this study were maintained in a growth cabinet set to 16 h of light at 25°C and 8 h of darkness at 23°C, with 70% relative humidity. An infectious cDNA clone of ANRSV-ZYZ (pRS) or its derivative (GFP-tagged clone, pRS-G) (Wang et al., 2021) was served as the backbone to develop the indicated recombinant clones in this study. For sap rub-inoculation assays, *N. benthamiana* plants were respectively inoculated with pPasFru-G (Gou et al., 2023), pHNu-GFP (Hu et al., 2020) and pCBTuMV-GFP/mCherry (Dai et al., 2021), and the infected leaf tissues were used to prepare the homogenates containing GFP-tagged TelMV (TelMV-GFP), PVMV (PVMV-GFP) or TuMV (TuMV-GFP).

### Construction of ANRSV-derived cDNA clones

Either pRS-G or pRS-G was used as the backbone to construct a series of ANRSV-derived virus clones, including pRS-G(△HCPro1), pRS-G(△HCPro2), pRS-G(tuHCPro), pRS-G(tbP19), pRS-GFP-HCPro2, pRS-G-2×Strep-HCPro2 and pRS-G(△HCPro1)-2×Strep-HCPro2. These clones were constructed by a similar strategy, mainly based on standard DNA manipulation technologies including overlapping PCR. Herein, the detailed description for the creation of pRS-G(△HCPro2), in which the complete HCPro2-coding sequence in ANRSV was deleted, was exemplified. Two PCR reactions with pRS-G as the template were performed using corresponding primer sets PCB301-F/SOE-HP2-R and SOE-HP2-F/RSV-3-R (Supplemental Table S2). A mixture of resulting PCR products was used as the template for overlapping PCR with the primer set PCB301-F/ RSV-3-R (Supplemental Table S2). The obtained fragment was inserted back into pRS-G by using *Pme* I / *Mlu* I sites to generate pRS-G(△HCPro2). pRS-G(△GDD), a replication-defective virus clone, was created via the removal of strictly-conserved GDD motif in viral RNA polymerase (NIb). Two fragments upstream and downstream of GDD motif in pRS-G were amplified with corresponding primer sets RSV-5-F/SOE-GDD-R and SOE-GDD-F/RSV-5-R (Supplemental Table S2), and then mixed as the template for overlapping PCR with primer set RSV-5-F/RSV-5-R (Supplemental Table S2). The obtained fragment was inserted back into *EcoR* I/*Sal* I-treated pRS-G to generate pRS-G(△GDD).

### Construction of binary plant-expression vectors

For RSS assay, four plasmids, including pCaM-HCPro1-HCPro2-HA, pCaM-HCPro1-HA, pCaM-HCPro2-HA, and pCaM-ssHCPro2-HA, were constructed for respective expression of HCPro1-HCPro2-HA, HCPro1-HA, HCPro2-HA and ssHCPro2-HA. The coding regions of them were amplified from pRS-G (Wang et al., 2021) or pSS-I-G (Qin et al., 2021), and individually integrated into a binary plant expression vector pCaMterX (Joensuu et al., 2010) by using *Xho* I / *Kpn* I sites. GenBank accession numbers of *NbRbCS* and *NbFNR* sequences are QCS40508.1 and QAV53876.1, respectively. We referred to these sequences to design primer sets to amplify complete or partial sequences of *NbRbCS* and *NbFNR* from *N. benthamiana* (Supplementary Table S1). For TRV-based VIGS analysis, SGN VIGS Tool (https://vigs.solgenomics.net) was employed to design two pairs of primers TRV-NbRbCS-F/TRV-NbRbCS-R and TRV-NbFNR-F/TRV-NbFNR-R (Supplemental Table S1) for amplifying an approximately 300-bp fragment of either *NbRbCS* or *NbFNR*. The obtained fragments were individually cloned into pTRV2 (Liu, et al, 2002) by utility of *BamH* I / *Xho* I sites to obtain pTRV2-NbRbCS and pTRV2-NbFNR (Supplemental Table S1). For Y2H, BiFC and Co-IP assays, the corresponding plasmids were generated by using Gateway cloning technology. Briefly, the complete coding sequence of each tested cistron was engineered into the entry clone - pDONR221, and then transferred into the desired gateway-compatible destination vectors, including pGADT7-DEST, pGBKT7-DEST, pEarleygate201-YN, pEarleygate202-YC, or/and pBA-FLAG-4myc-DC (Earley et al., 2006; Lu et al., 2010; Zhu et al., 2011). In addition, we constructed four plasmids (pCaM-GFP-HCPro2, pCaM-CI-mCherry, pCaM-CP-mCherry, and pCaM-NbRbCS-mCherry) for respective expression of GFP-HCPro2, CI-mCherry, CP-mCherry, and NbRbCS-mCherry. For this, we firstly amplified complete *GFP* and *mCherry* sequences from pVPH-GFP//mCherry (Cui & Wang, 2016), individually engineered them to pCaMterX, and obtained two intermediate vectors - pCaM-GFP and pCaM-mCherry. Then, the complete coding sequences of HCPro2, CI, CP, and NbRbCS were individually integrated into pCaM-GFP or pCaM-mCherry to produce the four plasmids via seamless cloning or restriction endonuclease digestion - T4 DNA ligation strategy.

All plasmids in this study were verified by Sanger DNA sequencing.

### Agrobacterium infiltration and sap rub-inoculation

*Agrobacterium* (strain GV3101)-mediated transformation was performed in *N. benthamiana*, essentially as previously described (Qin et al., 2021; Wang et al., 2021). Fully expanded leaves were infiltrated with agrobacterial cultures harboring relevant plasmids. *N. benthamiana* seedlings at 3- to 5-leaf stage were used for infectivity test of ANRSV-derived cDNA clones. The seedlings at 6- to 8-leaf stage are used for transient expression of genes of interest. For TRV-VIGS assays, two agrobacterial cultures harboring pTRV1 along with pTRV2-GUS (TRV-GUS), pTRV2-NbRbCS or pTRV2-NbFNR were mixed in a ratio of 1:1 (final OD_600_ = 0.3 per culture), and then infiltrated into the seedlings at 3- to 5-leaf stage. A sap rub inoculation assay was employed to test the infectivity of ANRSV-GFP, PVMV-GFP, TelMV-GFP and TuMV-GFP in *RbCS*- or/and *FNR*-silenced *N. benthamiana* plants. Briefly, leaf tissues of *N. benthamiana* plants infected by ANRSV-GFP, PVMV-GFP, TelMV-GFP or TuMV-GFP were ground in phosphate buffer (KH_2_PO_4_, 0.01 M, Na_2_HPO_4_, 0.01 M; pH 6.8) in a ratio of 1:10 (gram per milliliter). The resulting homogenate as an inoculum source was immediately rub inoculated into the newly developed leaves of *N. benthamiana* plants pre-inoculated with TRV-GUS (as the control), TRV-NbRbCS, or TRV-NbFNR. One leaf per plant was rubbed twice from the leaf bottom to the top using an inoculum-dipped forefinger.

### Yeast two-hybrid assays

Y2H assays were performed according to Yeastmaker™ Yeast Transformation System 2 User Manual (Clontech). The coding sequences corresponding to HCPro1, HCPro2, P3, P3N-PIPO, 6K1, CI, 6K2, NIa-Pro, VPg, NIb, CP and NbRbCS were individually cloned into either pGBKT7-DEST for fusing with GAL4 DNA binding domain (BD), or pGADT7-DEST for fusing with GAL4 activation domain (AD). Subsequently, yeast competent cells (Y2H gold) (WEIDI) was co-transformed with various combinations of bait and prey constructs, followed by 10-fold serial dilution and plating onto either the synthetic defined (SD) yeast leucine and tryptophan dropout medium (SD/-Leu/-Trp) or the leucine, tryptophan, histidine and adenine dropout medium (SD/-Leu/-Trp/-His/-Ade). The transformants were allowed by 4- to 6-day growth on the dropout mediums at 28 °C.

### Bimolecular fluorescence complementation

For BiFC assays, the coding sequences corresponding to HCPro2, CI, CP, P3N-PIPO, NbRbCS, and the N-terminal (N2) and C-terminal domains (D2) of HCPro2 were individually integrated into p35S-gatewayYN or/and p35S-gatewayYC (Lu et al., 2010) for the expression of these proteins fused with N-terminal half (YN) or C-terminal half of YFP (YC). Two agrobacterial cultures harboring the combination of YN- and YC-constructs were mixed in a ratio of 1:1 (final OD_600_ = 0.3 per culture), and then infiltrated into the fully developed leaves of *N. benthamiana*. The inoculated leaves were examined by an inverted fluorescence microscope (BX53F, OLYMPUS) or confocal microscopy at the indicated time points.

### Co-immunoprecipitation

Total proteins were extracted from 1 g of the co-inoculated leaves of *N. benthamiana* by using 2 mL of ice-cold immunoprecipitation buffer (10% [v/v] glycerol, 25 mM Tris-HCI, pH 7.5, 150 mM NaCl, 10 mM DTT, 1 mM EDTA, 1 × Protease Inhibitor Cocktail, For Plant Cell (Sangon Biotech), and 0.15% [v/v] Nonidet P-40). Protein extracts were incubated with GFP-Trap (ChromoTek) beads for 1h at 4°C. The beads were collected and washed with the buffer (10 mM Tris-HCl pH 7.5, 150 mM NaCl, 0.05 % Nonidet™ P40 Substitute, 0.5 mM EDTA). Total protein extracts prior to (Input) and after immunoprecipitation (IP) were analysed by immunoblotting using anti-GFP and anti-Myc polyclonal antibodies (Abcam), essentially as previously described (Qin et al., 2021).

### Streptavidin affinity purification and liquid chromatography tandem mass spectrometry

Streptavidin affinity purification in combination with LC-MS/MS were essentially performed following a previously described protocol (Hu et al., 2023). In brief, the upper non-inoculated leaves were collected from *N. benthamiana* plants infiltrated with pRS-G-2×Strep-HCPro2, pRS-G(ΔHCPro1)-2×Strep-HCPro2 or pRS-G at 12 dpi. The leaf tissues (30 g / treatment) were fine grinded, following by the addition of nondenaturing lysis buffer. The homogenate was centrifuged to produce a supernatant. The supernatant with an addition of Avidin was subjected to streptavidin affinity purification by using a Strep-Taction affinity column, followed by column-washing (five times). The obtained products were analyzed by either SDS-PAGE or immunoblotting with anti-Strep-tag II monoclonal antibody (BBI), followed by LC-MS/MS analysis (Beijing Bio-Tech Pack Technology Company Ltd.). The resulting data were analyzed by MaxQuant software to search against UniProt protein database (https://www.uniprot.org/) to identify the candidate proteins.

### Subcellular fractionation assay

We referred to several previous documents (Han& Sanfaçon, 2003; Gardiner & Chrispeels, 1975; Schaad et al., 1997) to perform subcellular fractionation assay. The leaf tissues (1 g per treatment) were collected from *N. benthamiana* plants inoculated with pRS-G or pRS-GFP-HCPro2, and fine homogenized in 4 mL of lysis buffer (50 mM Tris-HCl, pH 7.4, 15 mM MgCl_2_, 10 mM KCl, 20% glycerol, 1 × Protease Inhibitor Cocktail, For Plant Cell). The homogenate was centrifuged at 1000 *g* for 5 min at 4°C to remove the debris, and the supernatants (S1) was obtained. The S1 was centrifuged at 3700 *g* for 10 min at 4°C, resulting in supernatant (S3) and crude pellet (P3) fractions. The P3 fraction includes nuclei, chloroplasts and cell wall. Subsequently, the S3 was centrifuged at 30000 *g* for 50 min at 4°C to separate the soluble (S30) and crude membrane (P30) fractions, and then the P30 pellet was resuspended in the extraction buffer.

### RNA extraction, Northern blot and real-time RT-qPCR

Total RNAs were extracted from *N. benthamiana* leaves by using TRNzol Universal Reagent (TIANGEN). Norther blot was employed to detect *GFP* mRNA accumulation. The procedures, including total RNAs separation, transfer and fixation into nylon membrane, and hybridization with *GFP*-specific probes, were exactly the same with our previous description (Qin et al., 2021; Hu et al., 2023). The hybridization signals in the blotted membranes were detected with the substrates of enhanced chemiluminescence detection reagents (Thermo Fisher Scientific), visualized using an ImageQuant LAS 4000 mini-biomolecular imager (GE Healthcare). We performed real-time RT-qPCR to relatively quantity either viral genomic RNAs or endogenous gene transcripts. In the assays, an amount of 1 μg of total RNAs per biological sample was treated with DNase I (Thermo Scientific), and subsequently reverse-transcribed into cDNAs using RevertAid First Strand cDNA Synthesis Kit (Thermo Scientific) with random hexamer primers. The qPCR reactions with SuperReal PreMix Plus (SYBR green) (Tiangen) were carried out with Applied Biosystems QuantStudio 5 (Thermo Scientific) following the manufacturer’s instructions. The primers used in qPCR were listed in Supplemental Table S1. *NbActin* transcripts were determined to normalize the data.

### Aniline blue staining

Aniline blue solution is prepared before use via mixing 0.1% aniline blue (Sigma-Aldrich)-water solution and 1 M glycerol solution (in water) in a ratio of 2:3. The mixture was infiltrated into *N. benthamiana* leaves by using a 1 mL needle-free syringe. Thirty minutes later, aniline blue fluorescence was observed under confocal microscope using DAPI diode laser for excitation and filtering the emission.

### Confocal microscopy

The epidermal cells of inoculated leaves with relevant plasmids were observed under a confocal microscopy (FV1000, OLYMPUS) with a 20×water immersion objective. Excitation wavelengths and emission filters were 488 nm/bandpass 500-530 nm for GFP or YFP, and 543 nm/bandpass 580-620 nm for mCherry, 405 nm/band-pass 442-472 nm for aniline blue fluorochrome.

## Acknowledgments

This work is supported by grants from the National Natural Science Foundation of China (32360651, 32060603, 32372484) and Collaborative Innovation Center of Nanfan and High-Efficiency Tropical Agriculture, Hainan University (XTCX2022NYB11). We thank Prof. Jingsheng Xu (Fujian Agriculture and Forestry University) for the assistance in aniline blue staining, and Dr. Aiming Wang (Agriculture and Agri-Food Canada) and Dr. Guanwei Wu (Ningbo University) for critical suggestions. HC, ZD and LQ conceived the project. LQ and HL carried out experiments. HC supervised the work. All authors analyzed and discussed the data. HC and LQ draft the manuscript, and the remaining authors revise it. All authors reviewed and approved the manuscript. We declare no conflicts of interest.

## References

Anandalakshmi, R., Pruss, G. J., Ge, X., Marathe, R., Mallory, A. C., Smith, T. H., Vance, V. B. 1998. A viral suppressor of gene silencing in plants. Proceedings of the National Academy of Sciences, 95, 13079–13084.

Bar-On, Y. M., Milo, R. 2019. The global mass and average rate of rubisco. Proceedings of the National Academy of Sciences, 116, 4738–4743.

Beauchemin, C., Laliberté, J. F. 2007. The poly (A) binding protein is internalized in virus-induced vesicles or redistributed to the nucleolus during turnip mosaic virus infection. Journal of Virology, 81, 10905–10913.

Bhattacharyya, D., Chakraborty, S. 2018. Chloroplast: the Trojan horse in plant–virus interaction. Molecular Plant Pathology, 19, 504–518.

Bracher, A., Whitney, S. M., Hartl, F. U., Hayer-Hartl, M. 2017. Biogenesis and metabolic maintenance of Rubisco. Annual Review of Plant Biology, 68, 29–60.

Butkovic, A., Dolja, V. V., Koonin, E. V., Krupovic, M. 2023. Plant virus movement proteins originated from jelly-roll capsid proteins. PLoS Biology, 21, e3002157.

Carrington, J. C., Jensen, P. E., Schaad, M. C. 1998. Genetic evidence for an essential role for potyvirus CI protein in cell-to-cell movement. The Plant Journal, 14, 393–400.

Chen, I. H., Chen, X. Y., Chiu, G. Z., Huang, Y. P., Hsu, Y. H., Tsai, C. H. 2022. The function of chloroplast ferredoxin-NADP+ oxidoreductase positively regulates the accumulation of bamboo mosaic virus in *Nicotiana benthamiana*. Molecular Plant Pathology, 23, 503–515.

Cheng, D. J., Xu, X. J., Yan, Z. Y., Tettey, C. K., Fang, L., Yang, G. L., …, Li, X. D. 2021. The chloroplast ribosomal protein large subunit 1 interacts with viral polymerase and promotes virus infection. Plant Physiology, 187, 174–186.

Cheng, G., Dong, M., Xu, Q., Peng, L., Yang, Z., Wei, T., Xu, J. 2017. Dissecting the molecular mechanism of the subcellular localization and cell-to-cell movement of the sugarcane mosaic virus P3N-PIPO. Scientific Reports, 7, 9868.

Cheng, G., Yang, Z., Zhang, H., Zhang, J., & Xu, J. (2020). Remorin interacting with PCaP1 impairs turnip mosaic virus intercellular movement but is antagonised by VPg. New Phytologist, 225(5), 2122–2139.

Choi, I. R., Horken, K. M., Stenger, D. C., French, R. 2005. An internal RNA element in the P3 cistron of wheat streak mosaic virus revealed by synonymous mutations that affect both movement and replication. Journal of General Virology, 86, 2605–2614.

Chung, B. Y. W., Miller, W. A., Atkins, J. F., Firth, A. E. 2008. An overlapping essential gene in the *Potyviridae*. Proceedings of the National Academy of Sciences, 105, 5897–5902.

Cui, H., Wang, A. 2019. The biological impact of the hypervariable N-terminal region of potyviral genomes. Annual Review of Virology, 6, 255–274.

Cui, X., Yaghmaiean, H., Wu, G., Wu, X., Chen, X., Thorn, G., Wang, A. 2017. The C-terminal region of the turnip mosaic virus P3 protein is essential for viral infection via targeting P3 to the viral replication complex. Virology, 510, 147–155.

Dai, Z., He, R., Bernards, M. A., Wang, A. 2020. The cis-expression of the coat protein of turnip mosaic virus is essential for viral intercellular movement in plants. Molecular Plant Pathology, 21, 1194–1211.

De, S., Pollari, M., Varjosalo, M., Mäkinen, K. 2020. Association of host protein VARICOSE with HCPro within a multiprotein complex is crucial for RNA silencing suppression, translation, encapsidation and systemic spread of potato virus A infection. PLoS Pathogens, 16, e1008956.

Deng, P., Wu, Z., Wang, A. 2015. The multifunctional protein CI of potyviruses plays interlinked and distinct roles in viral genome replication and intercellular movement. Virology Journal, 12, 1–11.

Dolja, V. V., Haldeman-Cahill, R., Montgomery, A. E., Vandenbosch, K. A., Carrington, J. C. 1995. Capsid protein determinants involved in cell-to-cell and long distance movement of tobacco etch potyvirus. Virology, 206, 1007–1016.

Dolja, V. V., Haldeman, R., Robertson, N. L., Dougherty, W. G., Carrington, J. C. 1994. Distinct functions of capsid protein in assembly and movement of tobacco etch potyvirus in plants. The EMBO Journal, 13, 1482–1491.

Earley, K. W., Haag, J. R., Pontes, O., Opper, K., Juehne, T., Song, K., Pikaard, C. S. 2006. Gateway-compatible vectors for plant functional genomics and proteomics. The Plant Journal, 45, 616–629.

Faulkner, C. 2018. Plasmodesmata and the symplast. Current Biology, 28, R1374–R1378.

Feki, S., Loukili, M. J., Triki-Marrakchi, R., Karimova, G., Old, I., Ounouna, H., …, Elgaaied, A. B. A. 2005. Interaction between tobacco ribulose-l, 5-biphosphate carboxylase/oxygenase large subunit (RubisCO-LSU) and the PVY coat protein (PVY-CP). European Journal of Plant Pathology, 112, 221–234.

Gabrenaite-Verkhovskaya, R., Andreev, I. A., Kalinina, N. O., Torrance, L., Taliansky, M. E., Mäkinen, K. 2008. Cylindrical inclusion protein of potato virus A is associated with a subpopulation of particles isolated from infected plants. Journal of General Virology, 89, 829–838.

Gardiner, M., Chrispeels, M. J. 1975. Involvement of the Golgi apparatus in the synthesis and secretion of hydroxyproline-rich cell wall glycoproteins. Plant Physiology, 55, 536–541.

Geng, C., Cong, Q. Q., Li, X. D., Mou, A. L., Gao, R., Liu, J. L., Tian, Y. P. 2015. Developmentally regulated plasma membrane protein of *Nicotiana benthamiana* contributes to potyvirus movement and transports to plasmodesmata via the early secretory pathway and the actomyosin system. Plant Physiology, 167, 394–410.

Gou, B., Dai, Z., Qin, L., Wang, Y., Liu, H., Wang, L., …, Cui, H. 2023. A zinc finger motif in the P1 N terminus, highly conserved in a subset of potyviruses, is associated with the host range and fitness of telosma mosaic virus. Journal of Virology, 97, e01444–22.

Grimm, R., Grimm, M., Eckerskorn, C., Pohlmeyer, K., Röhl, T., Soll, J. 1997. Postimport methylation of the small subunit of ribulose-1, 5-bisphosphate carboxylase in chloroplasts. FEBS Letters, 408, 350–354.

Guo, D., Merits, A., Saarma, M. 1999. Self-association and mapping of interaction domains of helper component-proteinase of potato A potyvirus. Journal of General Virology, 80, 1127–1131.

Guo, D., Rajamäki, M. L., Saarma, M., Valkonen, J. P. 2001. Towards a protein interaction map of potyviruses: protein interaction matrixes of two potyviruses based on the yeast two-hybrid system. Journal of General Virology, 82, 935–939.

Hafrén, A., Lõhmus, A., Mäkinen, K. 2015. Formation of potato virus A-induced RNA granules and viral translation are interrelated processes required for optimal virus accumulation. PLoS Pathogens, 11, e1005314.

Han, K., Zheng, H., Yan, D., Zhou, H., Jia, Z., Zhai, Y., …, Yan, F. 2023. Pepper mild mottle virus coat protein interacts with pepper chloroplast outer envelope membrane protein OMP24 to inhibit antiviral immunity in plants. Horticulture Research, 10, uhad046.

Han, S., Sanfaçon, H. 2003. Tomato ringspot virus proteins containing the nucleoside triphosphate binding domain are transmembrane proteins that associate with the endoplasmic reticulum and cofractionate with replication complexes. Journal of Virology, 77, 523–534.

Harries, P., Ding, B. 2011. Cellular factors in plant virus movement: at the leading edge of macromolecular trafficking in plants. Virology, 411, 237–243.

Heinlein, M. 2015. Plasmodesmata: channels for viruses on the move. Plasmodesmata: Methods and Protocols, 25–52.

Hu, W., Dai, Z., Liu, P., Deng, C., Shen, W., Li, Z., Cui, H. 2023. The single distinct leader protease encoded by alpinia oxyphylla mosaic virus (genus *Macluravirus*) suppresses RNA silencing through interfering with double-stranded RNA synthesis. Phytopathology, 113, 1103–1114.

Hu, W., Qin, L., Yan, H., Miao, W., Cui, H., Liu, W. 2020. Use of an infectious cDNA clone of pepper veinal mottle virus to confirm the etiology of a disease in *Capsicum chinense*. Phytopathology, 110, 80–84.

Inoue-Nagata, A. K., Jordan, R., Kreuze, J., Li, F., López-Moya, J. J., Mäkinen, K., …, ICTV Report Consortium. 2022. ICTV virus taxonomy profile: *Potyviridae* 2022. Journal of General Virology, 103, 001738.

Jackson, A. O., Lim, H. S., Bragg, J., Ganesan, U., Lee, M. Y. 2009. Hordeivirus replication, movement, and pathogenesis. Annual Review of Phytopathology, 47, 385–422.

Ji, M., Zhao, J., Han, K., Cui, W., Wu, X., Chen, B., …, Yan, F. 2021. Turnip mosaic virus P1 suppresses JA biosynthesis by degrading cpSRP54 that delivers AOCs onto the thylakoid membrane to facilitate viral infection. PLoS Pathogens, 17, e1010108.

Joensuu, J. J., Conley, A. J., Lienemann, M., Brandle, J. E., Linder, M. B., Menassa, R. 2010. Hydrophobin fusions for high-level transient protein expression and purification in *Nicotiana benthamiana*. Plant Physiology, 152, 622–633.

Kang, S. H., Lim, W. S., Kim, K. H. 2004. A protein interaction map of soybean mosaic virus strain G7H based on the yeast two-hybrid system. Molecules & Cells, 18, 122–126.

Kasschau, K. D., Carrington, J. C. 1998. A counterdefensive strategy of plant viruses: suppression of posttranscriptional gene silencing. Cell, 95, 461–470.

Kawakami, S., Watanabe, Y., Beachy, R. N. 2004. Tobacco mosaic virus infection spreads cell to cell as intact replication complexes. Proceedings of the National Academy of Sciences, 101, 6291–6296.

Kumar, S., Karmakar, R., Gupta, I., Patel, A. K. 2020. Interaction of potyvirus helper component-proteinase (HcPro) with RuBisCO and nucleosome in viral infections of plants. Plant Physiology and Biochemistry, 151, 313–322.

Langenberg, W. G., Purcifull, D. E. 1989. Interactions between pepper ringspot virus and cylindrical inclusions of two potyviruses. Journal of Ultrastructure and Molecular Structure Research, 102, 53–58.

Langenberg, W. G. 1991. Cylindrical inclusion bodies of wheat streak mosaic virus and three other potyviruses only self-assemble in mixed infections. Journal of General Virology, 72, 493–497.

Langenberg, W. G. 1993. Structural proteins of three viruses in the *Potyviridae* adhere only to their homologous cylindrical inclusions in mixed infections. Journal of Structural Biology, 110, 188–195.

Laporte, C., Vetter, G., Loudes, A. M., Robinson, D. G., Hillmer, S., Stussi-Garaud, C., Ritzenthaler, C. 2003. Involvement of the secretory pathway and the cytoskeleton in intracellular targeting and tubule assembly of grapevine fanleaf virus movement protein in tobacco BY-2 cells. The Plant Cell, 15, 2058–2075.

Li, F., Huang, C., Li, Z., Zhou, X. 2014. Suppression of RNA silencing by a plant DNA virus satellite requires a host calmodulin-like protein to repress RDR6 expression. PLoS Pathogens, 10, e1003921.

Li, Y., Cui, H., Cui, X., Wang, A. 2016. The altered photosynthetic machinery during compatible virus infection. Current Opinion in Virology, 17, 19–24.

Li, Z., Liu, S. L., Montes-Serey, C., Walley, J. W., Aung, K. 2022. Plasmodesmata-located proteins regulate plasmodesmal function at specific cell interfaces in Arabidopsis. BioRxiv, 2022–08

Lim, H. S., Bragg, J. N., Ganesan, U., Ruzin, S., Schichnes, D., Lee, M. Y., …, Jackson, A. O. 2009. Subcellular localization of the barley stripe mosaic virus triple gene block proteins. Journal of Virology, 83, 9432–9448.

Lin, L., Luo, Z., Yan, F., Lu, Y., Zheng, H., Chen, J. 2011. Interaction between potyvirus P3 and ribulose-1, 5-bisphosphate carboxylase/oxygenase (RubisCO) of host plants. Virus Genes, 43, 90–92.

Liu, Y., Schiff, M., Marathe, R., Dinesh-Kumar, S. P. 2002. Tobacco Rar1, EDS1 and NPR1/NIM1 like genes are required for N-mediated resistance to tobacco mosaic virus. The Plant Journal, 30, 415–429.

Lõhmus, A., Varjosalo, M., Mäkinen, K. 2016. Protein composition of 6K2-induced membrane structures formed during potato virus A infection. Molecular Plant Pathology, 17, 943–958.

López, L., Urzainqui, A., Domínguez, E., García, J. A. 2001. Identification of an N-terminal domain of the plum pox potyvirus CI RNA helicase involved in self-interaction in a yeast two-hybrid system. Journal of General Virology, 82, 677–686.

Lu, Q., Tang, X., Tian, G., Wang, F., Liu, K., Nguyen, V. I., …, Cui, Y. 2010. Arabidopsis homolog of the yeast TREX-2 mRNA export complex: components and anchoring nucleoporin. The Plant Journal, 61, 259–270.

Lucas, W. J., Ham, B. K., Kim, J. Y. 2009. Plasmodesmata–bridging the gap between neighboring plant cells. Trends in Cell Biology, 19, 495–503.

Mao, Y., Catherall, E., Díaz-Ramos, A., Greiff, G. R., Azinas, S., Gunn, L., McCormick, A. J. 2023. The small subunit of Rubisco and its potential as an engineering target. Journal of Experimental Botany, 74, 543–561.

Martínez-Turiño, S., García, J. A. 2020. Potyviral coat protein and genomic RNA: a striking partnership leading virion assembly and more. Advances in Virus Research, 108, 165–211.

Medina-Puche, L., Tan, H., Dogra, V., Wu, M., Rosas-Diaz, T., Wang, L., …, Lozano-Duran, R. 2020. A defense pathway linking plasma membrane and chloroplasts and co-opted by pathogens. Cell, 182, 1109–1124.

Mingot, A., Valli, A., Rodamilans, B., San León, D., Baulcombe, D. C., García, J. A., López-Moya, J. J. 2016. The P1N-PISPO trans-frame gene of sweet potato feathery mottle potyvirus is produced during virus infection and functions as an RNA silencing suppressor. Journal of Virology, 90, 3543–3557.

Navarro, J. A., Sanchez-Navarro, J. A., Pallas, V. 2019. Key checkpoints in the movement of plant viruses through the host. Advances in Virus Research, 104, 1–64.

Olspert, A., Chung, B. Y. W., Atkins, J. F., Carr, J. P., Firth, A. E. 2015. Transcriptional slippage in the positive-sense RNA virus family *Potyviridae*. EMBO Reports, 16, 995–1004.

Oparka, K. J. 2004. Getting the message across: how do plant cells exchange macromolecular complexes? Trends in Plant Science, 9, 33–41.

Otulak, K., Garbaczewska, G. 2012. Cytopathological potato virus Y structures during Solanaceous plants infection. Micron, 43, 839–850.

Park, S. H., Li, F., Renaud, J., Shen, W., Li, Y., Guo, L., …, Wang, A. 2017. NbEXPA1, an α-expansin, is plasmodesmata-specific and a novel host factor for potyviral infection. The Plant Journal, 92, 846–861.

Pasin, F., Daròs, J. A., Tzanetakis, I. E. 2022. Proteome expansion in the *Potyviridae* evolutionary radiation. FEMS Microbiology Reviews, 46, fuac011.

Pasin, F., Simón-Mateo, C., García, J. A. 2014. The hypervariable amino-terminus of P1 protease modulates potyviral replication and host defense responses. PLoS Pathogens, 10, e1003985.

Peng, Y. H., Kadoury, D., Gal-On, A., Huet, H., Wang, Y., Raccah, B. 1998. Mutations in the HC-Pro gene of zucchini yellow mosaic potyvirus: effects on aphid transmission and binding to purified virions. Journal of General Virology, 79, 897–904.

Pollari, M., De, S., Wang, A., Mäkinen, K. 2020. The potyviral silencing suppressor HCPro recruits and employs host ARGONAUTE1 in pro-viral functions. PLoS Pathogens, 16, e1008965.

Pouwels, J., Van Der Velden, T., Willemse, J., Borst, J. W., Van Lent, J., Bisseling, T., Wellink, J. 2004. Studies on the origin and structure of tubules made by the movement protein of cowpea mosaic virus. Journal of General Virology, 85, 3787–3796.

Prywes, N., Phillips, N. R., Tuck, O. T., Valentin-Alvarado, L. E., Savage, D. F. 2023. Rubisco function, evolution, and engineering. Annual Review of Biochemistry, 92.

Qin, L., Shen, W., Tang, Z., Hu, W., Shangguan, L., Wang, Y., …, Cui, H. 2020. A newly identified virus in the family *Potyviridae* encodes two leader cysteine proteases in tandem that evolved contrasting RNA silencing suppression functions. Journal of Virology, 95, 10–1128.

Reagan, B. C., Burch-Smith, T. M. 2020. Viruses reveal the secrets of plasmodesmal cell biology. Molecular Plant-Microbe Interactions, 33, 26–39.

Revers, F., García, J. A. 2015. Molecular biology of potyviruses. Advances in Virus Research, 92, 101–199.

Roberts, I. M., Wang, D., Findlay, K., Maule, A. J. 1998. Ultrastructural and temporal observations of the potyvirus cylindrical inclusions (CIs) show that the CI protein acts transiently in aiding virus movement. Virology, 245, 173–181.

Rodamilans, B., Valli, A., Mingot, A., San León, D., Baulcombe, D., López-Moya, J. J., García, J. A. 2015. RNA polymerase slippage as a mechanism for the production of frameshift gene products in plant viruses of the *Potyviridae* family. Journal of Virology, 89, 6965–6967.

Rodríguez-Cerezo, E., Findlay, K., Shaw, J. G., Lomonossoff, G. P., Qiu, S. G., Linstead, P., …, Risco, C. 1997. The coat and cylindrical inclusion proteins of a potyvirus are associated with connections between plant cells. Virology, 236, 296–306.

Rojas, M. R., Zerbini, F. M., Allison, R. F., Gilbertson, R. L., Lucas, W. J. 1997. Capsid protein and helper component-proteinase function as potyvirus cell-to-cell movement proteins. Virology, 237, 283–295.

Roudet-Tavert, G., German-Retana, S., Delaunay, T., Delécolle, B., Candresse, T., Le Gall, O. 2002. Interaction between potyvirus helper component-proteinase and capsid protein in infected plants. Journal of General Virology, 83, 1765–1770.

Sager, R. E., Lee, J. Y. 2018. Plasmodesmata at a glance. Journal of Cell Science, 131, jcs209346.

Saha, S., Lõhmus, A., Dutta, P., Pollari, M., Mäkinen, K. 2022. Interplay of HCPro and CP in the regulation of potato virus A RNA expression and encapsidation. Viruses, 14, 1233.

Schaad, M. C., Jensen, P. E., Carrington, J. C. 1997. Formation of plant RNA virus replication complexes on membranes: role of an endoplasmic reticulum-targeted viral protein. The EMBO Journal, 16, 4049–4059.

Schneider, C. A., Rasband, W. S., Eliceiri, K. W. 2012. NIH Image to ImageJ: 25 years of image analysis. Nature Methods, 9, 671–675.

Schoelz, J. E., Harries, P. A., Nelson, R. S. 2011. Intracellular transport of plant viruses: finding the door out of the cell. Molecular Plant, 4, 813–831.

Seo, J. K., Phan, M. S. V., Kang, S. H., Choi, H. S., Kim, K. H. 2013. The charged residues in the surface-exposed C-terminus of the soybean mosaic virus coat protein are critical for cell-to-cell movement. Virology, 446, 95–101.

Silhavy, D., Molnár, A., Lucioli, A., Szittya, G., Hornyik, C., Tavazza, M., Burgyán, J. 2002. A viral protein suppresses RNA silencing and binds silencing-generated, 21-to 25-nucleotide double-stranded RNAs. The EMBO Journal, 21, 3070–3080.

Stenger, D. C., French, R., Gildow, F. E. 2005. Complete deletion of wheat streak mosaic virus HC-Pro: a null mutant is viable for systemic infection. Journal of Virology, 79, 12077–12080.

Stenger, D. C., Hein, G. L., Gildow, F. E., Horken, K. M., French, R. 2005. Plant virus HC-Pro is a determinant of eriophyid mite transmission. Journal of Virology, 79, 9054–9061.

Sorel, M., García, J. A., German-Retana, S. 2014. The *Potyviridae* cylindrical inclusion helicase: a key multipartner and multifunctional protein. Molecular Plant-Microbe Interactions, 27, 215–226.

Torrance, L., Andreev, I. A., Gabrenaite-Verhovskaya, R., Cowan, G., Mäkinen, K., Taliansky, M. E. 2006. An unusual structure at one end of potato potyvirus particles. Journal of Molecular Biology, 357, 1–8.

Ueki, S., Citovsky, V. 2011. To gate, or not to gate: regulatory mechanisms for intercellular protein transport and virus movement in plants. Molecular Plant, 4, 782–793.

Untiveros, M., Olspert, A., Artola, K., Firth, A. E., Kreuze, J. F., Valkonen, J. P. 2016. A novel sweet potato potyvirus open reading frame (ORF) is expressed via polymerase slippage and suppresses RNA silencing. Molecular Plant Pathology, 17, 1111–1123.

Valli, A., Gallo, A., Calvo, M., Pérez, J. D. J., García, J. A. 2014. A novel role of the potyviral helper component proteinase contributes to enhance the yield of viral particles. Journal of Virology, 88, 9808–9818.

Valli, A. A., Gallo, A., Rodamilans, B., López-Moya, J. J., García, J. A. 2018. The HCPro from the *Potyviridae* family: an enviable multitasking helper component that every virus would like to have. Molecular Plant Pathology, 19, 744–763.

Verchot-Lubicz, J. 2005. A new cell-to-cell transport model for potexviruses. Molecular Plant-Microbe Interactions, 18, 283–290.

Verchot, J. 2022. Potato virus X: a global potato-infecting virus and type member of the *Potexvirus* genus. Molecular Plant Pathology, 23, 315–320.

Verchot-Lubicz, J., Torrance, L., Solovyev, A. G., Morozov, S. Y., Jackson, A. O., Gilmer, D. 2010. Varied movement strategies employed by triple gene block– encoding viruses. Molecular Plant-microbe Interactions, 23, 1231–1247.

Vijayapalani, P., Maeshima, M., Nagasaki-Takekuchi, N., Miller, W. A. 2012. Interaction of the trans-frame potyvirus protein P3N-PIPO with host protein PCaP1 facilitates potyvirus movement. PLoS Pathogens, 8, e1002639.

Wang, A. 2015. Dissecting the molecular network of virus-plant interactions: the complex roles of host factors. Annual Review of Phytopathology, 53, 45–66.

Wang, A. 2021. Cell-to-cell movement of plant viruses via plasmodesmata: a current perspective on potyviruses. Current Opinion in Virology, 48, 10–16.

Wang, Y., Shen, W., Dai, Z., Gou, B., Liu, H., Hu, W., …, Cui, H. 2021. Biological and molecular characterization of two closely related arepaviruses and their antagonistic interaction in *Nicotiana benthamiana*. Frontiers in Microbiology, 12, 755156.

Wei, T., Wang, A. 2008. Biogenesis of cytoplasmic membranous vesicles for plant potyvirus replication occurs at endoplasmic reticulum exit sites in a COPI-and COPII-dependent manner. Journal of Virology, 82, 12252–12264.

Wei, T., Zhang, C., Hong, J., Xiong, R., Kasschau, K. D., Zhou, X., …, Wang, A. 2010. Formation of complexes at plasmodesmata for potyvirus intercellular movement is mediated by the viral protein P3N-PIPO. PLoS Pathogens, 6, e1000962.

Wen, R. H., Hajimorad, M. R. 2010. Mutational analysis of the putative pipo of soybean mosaic virus suggests disruption of PIPO protein impedes movement. Virology, 400, 1–7.

Whitney, S. M., Houtz, R. L., Alonso, H. 2011. Advancing our understanding and capacity to engineer nature’s CO2-sequestering enzyme, Rubisco. Plant Physiology, 155, 27–35.

Wright, K. M., Wood, N. T., Roberts, A. G., Chapman, S., Boevink, P., MacKenzie, K. M., Oparka, K. J. 2007. Targeting of TMV movement protein to plasmodesmata requires the actin/ER network; evidence from FRAP. Traffic, 8, 21–31.

Wietrzynski, W., Traverso, E., Wollman, F. A., Wostrikoff, K. 2021. The state of oligomerization of Rubisco controls the rate of synthesis of the Rubisco large subunit in *Chlamydomonas reinhardtii*. The Plant Cell, 33, 1706–1727.

Wu, X., Cheng, X. 2020. Intercellular movement of plant RNA viruses: targeting replication complexes to the plasmodesma for both accuracy and efficiency. Traffic, 21, 725–736.

Yang, K., Ran, M., Li, Z., Hu, M., Zheng, L., Liu, W., …, Cui, H. 2018. Analysis of the complete genomic sequence of a novel virus, areca palm necrotic spindle-spot virus, reveals the existence of a new genus in the family *Potyviridae*. Archives of Virology, 163, 3471–3475.

Yang, K., Shen, W., Li, Y., Li, Z., Miao, W., Wang, A., Cui, H. 2019. Areca palm necrotic ringspot virus, classified within a recently proposed genus *Arepavirus* of the family *Potyviridae*, is associated with necrotic ringspot disease in areca palm. Phytopathology, 109, 887–894.

Yang, X., Li, Y., Wang, A. 2021. Research advances in potyviruses: from the laboratory bench to the field. Annual Review of Phytopathology, 59, 1–29.

You, Y., Shirako, Y. 2010. Bymovirus reverse genetics: requirements for RNA2-encoded proteins in systemic infection. Molecular Plant Pathology, 11, 383–394.

Zhao, J., Liu, Q., Zhang, H., Jia, Q., Hong, Y., Liu, Y. 2013. The rubisco small subunit is involved in tobamovirus movement and Tm-22-mediated extreme resistance. Plant Physiology, 161, 374–383.

Zhao, J., Zhang, X., Hong, Y., Liu, Y. 2016. Chloroplast in plant-virus interaction. Frontiers in Microbiology, 7, 1565.

Zhu, H., Hu, F., Wang, R., Zhou, X., Sze, S. H., Liou, L. W., …, Zhang, X. 2011. Arabidopsis Argonaute10 specifically sequesters miR166/165 to regulate shoot apical meristem development. Cell, 145, 242–256.

Zilian, E., Maiss, E. 2011. Detection of plum pox potyviral protein–protein interactions in planta using an optimized mRFP-based bimolecular fluorescence complementation system. Journal of General Virology, 92, 2711–2723.

